# Linker Histone H1.5 Contributes to Centromere Integrity in Human Cells

**DOI:** 10.1101/2025.06.03.657682

**Authors:** Ankita Saha, Minh Bui, Daniël P. Melters, Ganesan Arunkumar, Songjoon Baek, Reda S. Bentahar, Yamini Dalal

## Abstract

Mammalian H1 linker histones comprise a group of 11 non-allelic variants which have key roles in modulating chromatin. H1 variant specific genomic distribution contributes to fine tuning regulation of gene expression and chromatin architecture. Contradictory reports on the presence and role of H1 histones at centromeres led us to further investigate whether H1s impact centromeric chromatin. In this study, we focused on H1.5 and by *in vitro* assays we showed that H1.5 directly interacts with centromeric-protein A (CENP-A) mononucleosomes. Notably, our *in vitro* findings revealed that H1 variants H1.0 and H1.2 can also bind CENP-A nucleosomes, although with differing affinities and signatures, asserting centromeric localization may not be unique to H1.5. In human cells, H1.5 localized to the centromere and chromatin immuno-precipitation revealed an interaction between H1.5 with CENP-A nucleosomes. Knocking down of H1.5 resulted in the loss of centromeric α-satellite transcription, reduction in loading of new CENP-A, and the accumulation of mitotic defects. These data point to an unreported role for histone H1 in the regulation of mitotic integrity in human cells.

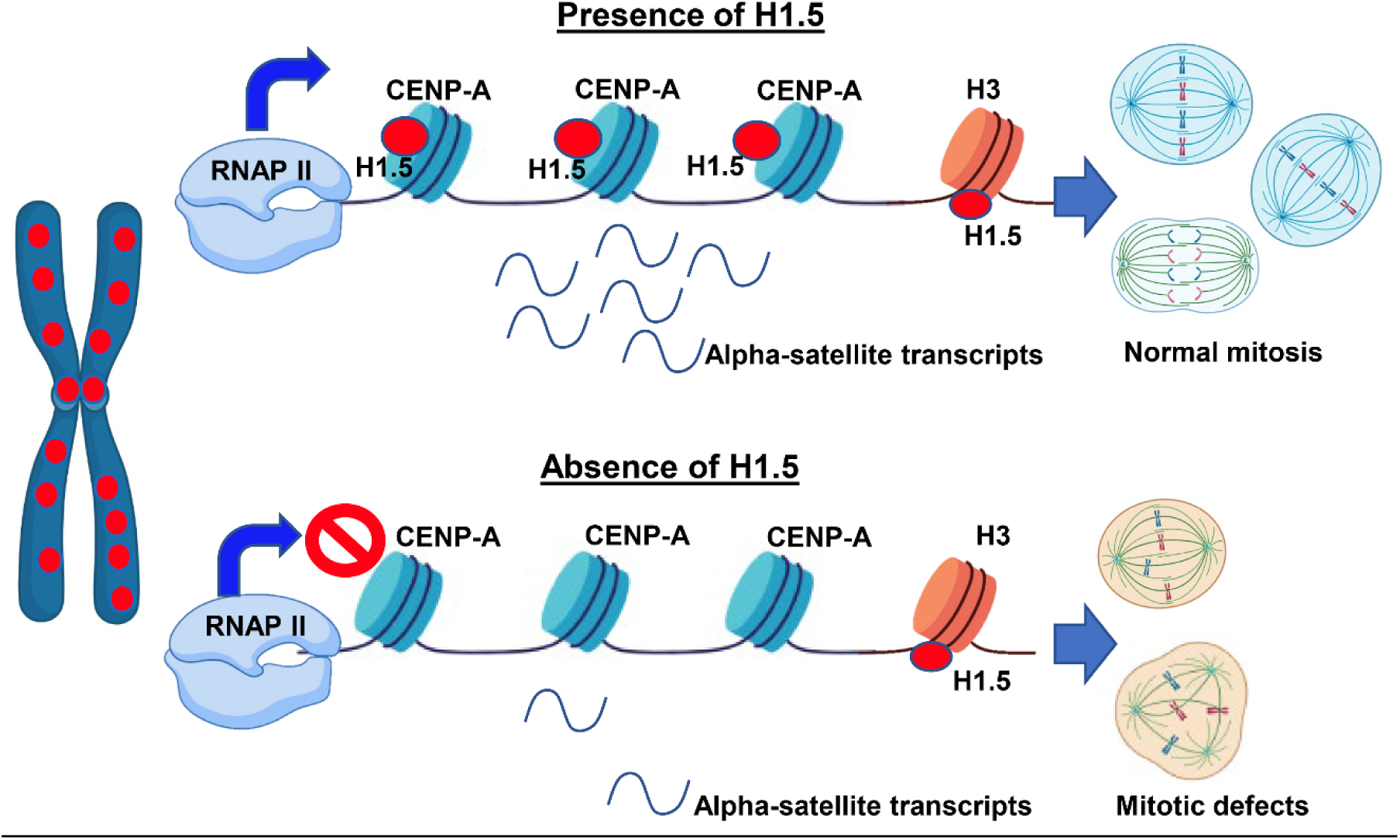
Graphical Abstract. Proposed model of H1.5 function at the centromere. When H1.5 is bound to the CENP-A nucleosomes at the centromere, the centromeric chromatin allows RNAPII based transcription to generate α-satellite transcripts, which in turn allows for *de novo* CENP-A loading. The absence of H1.5 blocks this transcription, leading to an accumulation of mitotic defects.

## INTRODUCTION

Chromatin is organized in a beads-on-a-string model, where every bead is a nucleosome that wraps 147 bp of DNA connected by linker DNA (1). The chromatin fiber is folded into higher-order structures, which are influenced greatly by the binding of the linker histone H1 protein. H1 binds nucleosomes at the nucleosome dyad, stabilizing the entry and exit DNA strands and permitting a more compact folding (2, 3). Folding and unfolding of the chromatin fiber is at the core of gene regulation (4, 5). *In vitro*, H1 compacts chromatin arrays into 30-nm fibers (6). *In vivo*, H1 is present in a stoichiometry of ∼0.5-to-1 per nucleosome (7, 8). Therefore, it is widely assumed that H1 has identical roles *in vivo* and *in vitro*, namely as a global transcriptional repressor (9, 10). There are 11 mammalian H1 variants, all of which have a tripartite structure with a central globular domain (GD), a short N-terminal domain (NTD) and a long, highly disordered C-terminal domain (CTD) (1–3, 11). The globular domain is highly conserved, whereas the NTD and CTD differ in composition and length. There are two widely described modes of H1-nucleosome binding, namely ‘on-dyad’ and ‘off-dyad’. The ‘on-dyad’ mode describes a central and symmetric positioning of the H1 GD, and terminal domains interact with both the nucleosome linker DNA arms. Meanwhile, ‘off-dyad’ is defined as the asymmetric positioning of the H1 GD adjacent to the dyad axis, where only one of the two linker DNA arms are in contact with the H1 terminal domains (12–16). The CTD of the *Xenopus* H1.0b dictate the precise positioning of the GD on the dyad and can regulate the H1-dependent asymmetric opening of nucleosomes, creating directionality along the DNA strand (17). With H1.0b CTD being able to engage with diverse DNA interfaces, it permits for tight regulation of nucleosome DNA unwrapping. It is therefore possible that each H1 variant could confer specific and unique functions in regulating access to nucleosome DNA and to higher order arrays (18–20). Indeed, work over the last couple decades showed that H1 plays a more nuanced role than simply serving as a transcriptional on/off switch. For example, knocking out either H1 genes in *Tetrahymena thermophila* only affected the expression of a few genes. H1.2 and H1.0 knockout studies in mammalian cells also showed only a subset of genes affected (21, 22).

Knockout studies of individual H1 genes in mice only had subtle phenotypes (23, 24) highlighting the ability of H1 variants to compensate for each other (19). Notably, a lethal phenotype was observed only when three H1 genes (H1.2, H1.3, and H1.4) were simultaneously knocked out (25). Knocking down of different H1s in human cells showed distinct phenotypes in H1 variant-specific manner, such as cell cycle arrest in H1.2 depleted cells or cell death upon H1.4 knock down. In contrast to the H1 knock-out mice, knocking down H1s in human cells did not result in expression compensation by other H1 variants (26). This was also observed in a different study where H1 depletion in Drosophila larvae led to only a modest upregulation of protein-coding genes and several downregulated genes. Surprisingly, transposable elements were strongly upregulated upon H1 depletion (27). These results suggest that while H1 variants may share some overlapping functions, they also possess variant-specific roles that are critical for regulating gene expression and cellular processes.

H1 variants are frequently misregulated in cancers. For example, the loss of the variant H1.0 has been identified as a prognostic biomarker for several cancers including malignant glioblastoma multiforme (GBM), breast cancer, melanoma, colorectal cancer, prostate adenocarcinoma, and bladder cancer (28). In follicular lymphoma, mutations in H1.4 and H1.5 were associated with altered chromatin states, downstream transformation of epigenetic states, and changes in gene expression (29). Linker histone H1.5 is also implicated in mammalian development, chromatin organization, and can serve as a valuable prognostic marker in cancer patients (30). These, along with other studies (31–36) point to an important role for H1 in maintaining genome integrity, and perturbations to H1 proteins or levels can contribute to cancer development (37, 38).

Chromosomal instability is a hallmark of many cancers, including GBM, and often arises from defects in centromere function and mitotic regulation (39, 40). The centromere is a specialized chromatin domain defined by the presence of histone H3 variant CENP-A, which replaces canonical H3 in nucleosomes at centromeric loci, and is essential for kinetochore assembly and accurate chromosome segregation (41–43). Although linker histones are known to compact chromatin and regulate gene expression, their role at the centromere has not been well defined. Partly, this is because of inconsistent results from the literature probing the interaction between H1 and CENP-A nucleosomes (44–46). Nearly 15 years ago, an *in vivo* fluorescence based (FRET) study demonstrated that H1s are at the centromere in close proximity to CENP-A, CENP-B, and CENP-C in human cells (45). By directly isolating centromeres from chromosomes, H1 was found at centromeres in midges (47). Strikingly, a cryoEM study reported little or no interaction between H1 and CENP-A and concluded that H1 may be excluded from centromeric chromatin for structural reasons (44). However, while our study work was under revision, an atomic force microscopy (AFM) paper concluded that H1.0 can bind CENP-A nucleosomes *in vitro* (48).

Thus, while the role of H1 in global chromatin compaction and transcriptional regulation is well established, its contribution to the centromere remains unresolved and is of obvious biological interest. Here, we use cell biology, biochemical, biophysical, and single molecule imaging approaches to meticulously dissect the role of a specific H1 variant at the normal human centromere using interdisciplinary approaches. First, we developed a novel native electromobility shift assay (EMSA) method to visualize *in vitro* reconstituted histone H1 onto either assembled H3- or CENP-A-mononucleosome that wrap a ∼200bp DNA fragment containing the 601-positioning sequence. We observed that CENP-A nucleosomes are directly associated with H1.0, H1.2, and H1.5 *in vitro*. We chose to focus on H1.5, as this linker histone is highly overexpressed in glioblastoma multiforme (GBM), and is a subject of ongoing work in the lab. We used *in vitro* biochemistry and AFM to study how H1.5 binds CENP-A. These data suggest that H1.5 can physically associate with CENP-A mononucleosomes, but in an unconventional binding mode, distinct from the known mechanism for interaction with canonical H3 nucleosomes. To examine a potential physiological relationship between H1.5 and CENP-A, we conducted experiments in astrocytes and HeLa cells. Immunofluorescence revealed H1.5 is enriched at the centromere and chromatin immunoprecipitation (ChIP) showed a physical interaction between H1.5 with CENP-A chromatin *in vivo*. Targeted depletion of H1.5 resulted in reduced transcription of non-coding centromeric α-satellite DNA and impaired *de novo* CENP-A loading. These disruptions were accompanied by the accumulation of mitotic defects, which could be partially rescued by reintroducing linker histone H1.5. Together, our results establish a role for linker histones in maintaining centromeric chromatin integrity and proper mitotic progression in human cells.

## MATERIAL AND METHODS

### Native EMSA in-gel Western Polyacrylamide Gel Electrophoresis (NEW-PAGE)

H3 or CENP-A nucleosomes were reconstituted by salt dialysis using a PCR-amplified 200 bp fragment containing the 601-positioning sequence with or without 5’-end labelled biotin, and with or without linker histone H1s. Reconstituted mononucleosomes were run under native conditions at 4°C (120 V for 2.5 hrs with 5% TBE acrylamide). Gel was stained with Streptavidin or GelStar for an hour (nucleic acid staining/GelStar signal is short-lived at <2 days), followed by overnight incubation with primary antibody against H1, CENP-A, or H3. Gel was imaged with LiCor M in gel mode with all detectable channels.

### MNase Protection Assay

After reconstitution, samples were MNase’ed for 4 min and quenched with EGTA, followed by proteinase K treatment. DNA samples were isolated with phenol-chloroform and electrophoresed in a 10% TBE acrylamide gel.

### Cell Culture and synchronization

Astrocyte (SVGp12) cells were cultured in EMEM medium supplemented with 10% fetal bovine serum (FBS), 5 mM glutamine, and 1 mM penicillin/streptomycin at 37°C in a humidified incubator with 5% CO2. For synchronization, SVGp12 cells were subjected to a double thymidine block protocol consisting of 5 mM thymidine (Sigma) for 22 hours, 12-hour release, a second thymidine block for 12 hours, followed by release and collection at 9 hours post-release. HeLa cells were cultured in DMEM, SW480 cells in RPMI-1640, and BJ fibroblasts in EMEM, all supplemented with 10% FBS and 1 mM penicillin/streptomycin at 37°C and 5% CO2.

### siH1.5 Transfections

For astrocytes, transient knockdowns were conducted using siRNA against H1.5 (Dharmacon: Cat# L-012049-02-0005) or siScramble ON-TARGETplus Non-targeting Control Pool (Dharmacon: Cat #D-001810-10-20) using Lipofectamine RNAimax (ThermoFisher Cat# 13778150) using manufacturer’s protocol and grown for 3 days before harvesting for downstream applications. H1.5 GFP plasmid (Addgene: Cata# 32898) transfection was carried out with Lipofectamine 3000 (ThermoFisher Cat # L3000015) using manufacturer’s protocol.

### Immuno-fluorescence and imaging

Cells were grown directly on glass coverslips and washed with PBS to remove residual media as previously described (49). In short, coverslips were permeabilized in lysis buffer (2.5 mM Tris-HCl pH 7.5, 0.5 mM NaCl, 1% Triton X-100, 0.4 M urea), then fixed in 4% paraformaldehyde in PBS.

After washing, cells were blocked in 0.5% BSA and 0.01% Triton X-100 in PBS for 30 minutes at room temperature. Cells were incubated overnight at 4°C with primary antibodies diluted in blocking buffer: rabbit anti-H1.5 (custom), goat anti-CENP-C (MBL International, Cat# PD030), and mouse anti-α-tubulin (Thermo Fisher Scientific, Cat# 62204; RRID:AB_1965960). Alexa Fluor-conjugated secondary antibodies (Invitrogen) were applied for 3 hours at room temperature.

DNA was counterstained with DAPI (50 ng/ml). Coverslips were mounted and imaged using a DeltaVision RT system (Applied Precision) fitted with a CoolSnap CCD camera on an Olympus IX70 microscope. Images were acquired at 0.1 μm z-sections and deconvolved using softWoRx software. Image processing and analysis were performed in ImageJ (NIH). Co-localization between H1.5 and CENP-C signals was assessed using the ‘Colocalization Finder’ plug-in within ImageJ.

### Metaphase spreads

Metaphase spreads were prepared as previously described (39). Briefly, SVGp12 cells were subjected to 8hrs of Colcemid treatment prior to harvest. The cells were subjected to hypotonic solution (75mM KCl), fixed with methanol: glacial acetic acid and dropped onto etched and chilled slides to obtain metaphase spreads. These prepared spreads were then dried or subject to immunofluorescence and then imaged.

### Chromatin immunoprecipitation

Chromatin immunoprecipitation (ChIP) was performed as previously outlined with modifications (50). Briefly, cells were harvested and washed with PBS and PBS containing 0.1% Tween 20.

Nuclei were released by incubation in TM2 buffer (20 mM Tris-HCl pH 8.0, 2 mM MgCl2) containing 0.5% Nonidet P-40 Substitute (Sigma Cat #74385). Nuclei were then washed with TM2 and digested with micrococcal nuclease (1 U MNase; Sigma Cat #N3755-500UN) for 8 minutes at 37°C in digestion buffer (10 mM Tris-HCl pH 8.0, 0.2 mM EDTA, 100 mM NaCl, 1.5 mM CaCl2). MNase digestion was quenched with 10 mM EGTA, and chromatin was extracted overnight at 4°C in 0.5X PBS containing protease inhibitor cocktail (Roche Cat #05056489001). The resulting mononucleosome-enriched chromatin was incubated with anti-CENP-A (Abcam Cat# ab13939) or anti-H1.5 (custom) antibodies and then eluted using Dynabeads Protein A (ThermoFisher Scientific Cat #10006D). Bead-bound complexes were washed 3 times with cold 0.5X PBS. Immunoprecipitated chromatin was either processed for Western blotting or DNA extraction followed by qRT-PCR.

### Quantitative real-time polymerase chain reaction

ChIP extracted DNA or cDNA samples were prepared using the PowerUP SYBR Green Master Mix (catalog no. A25777, Applied Biosystems, USA) following the manufacturer’s protocol. The quantitative reverse transcription PCR (qRT-PCR) was run on the StepOnePlus real-time PCR System (Applied Biosystems, USA), and relative quantification was performed using the 2^−ΔΔCT^ method. All reactions were performed in 20 μl volume and in triplicates on a 96-well plate (Applied biosystems). Melting curve analysis was performed for all the primer sets to check the specificity of the primers. Primer sequences used in this study are listed in Table 2.

### *In vitro* reconstitution and atomic force microscopy

H3 and CENP-A mononucleosomes (H3/H4 cat#16-0008, CENP-A/H4 cat#16-010, H2A/H2B cat#15-0311, Epicypher) were *in vitro* reconstituted on a 324-bp DNA fragment (PCR’ed from pGEM-3z601 plasmid from Addgene cat#26656) by stepwise salt-dialysis as described previously (50). The final buffer contained 150 mM NaCl, 2 mM MgCl2, 10 mM TRIS-Cl pH 8.0, and 1 mM EDTA. H1.5 was added to each sample and incubated for 1 hour at a ratio of nucleosome:H1.5 of 1:1. Samples were diluted 10-fold in 0.5x PBS + 2 mM MgCl2 buffer, incubated on an end-over-end rotator at RT for 30 minutes before depositing them on freshly cleaved V1 grade mica that was treated with aminopropyl-silantrane (APS) (51). Deposited samples were incubated on APS-functionalized mica for ∼10 minutes; excess buffer was rinsed with 2x 200 µL ultrapure, deionized water, and gently dried under an argon stream. Imaging was performed using standard AFM equipment (Asylum Research’s Cypher S AFM, Oxford Instruments) using silicon-nitride, oxide-sharpened probes (Bruker OTESPA-R3 with a nominal stiffness of 26 N/m and a nominal frequency of 300 kHz). Before each experiment, the spring constant of each cantilever was calibrated using the thermal noise method (51). Scan size was either 2x2 µm with a resolution of 1024 points and lines. Each reconstitution was performed in triplicate and at least 4 images per reconstitution were analyzed. For mononucleosomes the height and Feret’s diameter were measured to assess if H1.5 binds to individual H3 or CENP-A nucleosomes. Graphs were prepared using the ggplot2 package for R. Significance for nucleosome parameters were determined by the 2-sided Student t-test.

### SNAP-CENP-A construct, TMR Star, and data analysis

SNAP-tag was cloned upstream and in-frame to CENP-A with NheI and EcoRI restriction sites, and downstream of the CMV promoter. SNAP-CENP-A construct was transfected into HeLa cells using Neon Transfection System (ThermoFisher Scientific Cat #MPK5000) with 100 uL kit (ThermoFisher Scientific Cat #MPK10096), using the following parameters: 2 pulses of 1050 V/30 ms, and grown on coverslips with siH1.5 or siScramble.

Double thymidine block was done (see above) but during the first release, was removed and washed with PBS, and SNAP-Cell Block (NEB Cat #S9105S) was diluted 1:200 in 1 mL complete DMEM media, and cells incubated for 20 min. Coverslips were washed 3X with pre-warmed complete DMEM and released for the remainder of the 12-hr period. After the second release, cells were incubated for 30 min with SNAP-Cell TMR-star (NEB Cat #S9105S), washed 3X with pre-warm DMEM, released for the remainder of 14 hrs, and IF against endogenous CENP-A (AbCam Cat #ab13939) plus anti-mouse Alexa 488 (green). Imaging was done (see above) with FITC, TRITC, and DAPI channels. Images were captured by using a 100× objective at 0.1 μm z sections. Images were processed using softWoRx with deconvolution and projection. Two replicates were done.

The ImageJ ‘Colocalization’ plugin was used to determine whether SNAP-CENP-A colocalized with native CENP-A. Nuclei were considered to have colocalization if they had greater than 25% foci colocalizing (number of colocalized foci over the total number of native CENP-A foci).

### RNA Extraction and RNA-seq

Total RNA was extracted from SVGp12 cells following siScramble, siH1.0, or siH1.5 knockdown. Cells grown in T175 flasks were lysed directly in 1 ml of TRIzol reagent (Ambion, USA; Cat. No. 15596026) and incubated at room temperature for 5 minutes. Lysates were centrifuged at 12,000 rpm for 10 minutes at 4°C, and supernatants were transferred to fresh tubes. For phase separation, 200 µl of chloroform was added per 1 ml of TRIzol, followed by vortexing and incubation for 2 minutes at room temperature. After centrifugation (12,000 rpm, 15 minutes, 4°C), the aqueous phase was collected and mixed with 500 µl of isopropanol (Sigma-Aldrich; Cat. No. 534021) to precipitate RNA. RNA pellets were washed with cold 75% ethanol, air-dried, and resuspended in DEPC-treated ultrapure water (KD Medical, USA). Genomic DNA was removed by DNase I digestion (New England Biolabs, USA), and RNA quality and integrity were verified by 1% agarose gel electrophoresis containing GelStar nucleic acid stain (Lonza, USA; Cat. No. 50535). RNA samples were further purified by repeating TRIzol extraction and stored at −80°C.

For RNA expression profiling, RNA libraries were constructed using the Illumina TruSeq Stranded Total RNA Library Prep Kit (RS-122-2201) and sequenced on an Illumina NextSeq instrument using High 2.1 chemistry to generate 151 bp paired-end reads. Reads were trimmed for adapters and low-quality bases using Cutadapt (v1.18) and aligned to the human reference genome (GRCh38/hg38) with STAR/RSEM (v2.7.0f) using the parameters --alignIntronMin20 -- sjdbScore 1 --peOverlapNbasesMin10. Read counts were generated with Subread (v2.0.1) using Ensembl release 103 annotations. Differential gene expression was calculated using the DESeq2 package in RStudio (version 2024.09.0 Build 375, Posit Software). Statistical significance was defined at a false discovery rate–adjusted p-value of <0.01. Transcript abundance estimates were reported in units of FPKM (Fragments Per Kilobase of transcript per Million mapped reads). Data analysis and visualization were performed using R (version 4.4.0) and RStudio (version 2024.09.0 Build 375, Posit Software) with the following R/Bioconductor packages: DESeq2, ggplot2, pheatmap, UpSetR, clusterProfiler, and other standard tidyverse packages. All raw RNA-seq data have been deposited to the NCBI Gene Expression Omnibus (GEO) under accession number GSE242112.

### Statistical analyses

All statistical analyses were performed using GraphPad Prism software (v7.7e, GraphPad Software Inc., USA) or R (version 4.4.0, R Core Team) within RStudio (version 2024.09.0 Build 375, Posit Software, PBC). Data are presented as mean ± standard deviation (SD) or mean ± standard error of the mean (SEM) as indicated in figure legends. The number of independent biological replicates (n) is specified in the respective figure legends. Statistical significance between two groups was assessed using a two-tailed Student’s t-test or Mann–Whitney U test, as appropriate. For multiple group comparisons, two-way analysis of variance (ANOVA) was used with appropriate post hoc testing. A p-value less than 0.05 was considered statistically significant. p-values are presented in the figure legends as follows: *p < 0.05, **p < 0.01, ***p < 0.001, ****p < 0.0001. Illustrative diagrams were created using BioRender (BioRender, USA), Microsoft PowerPoint, and R.

### Acid Extraction of Histones

Histones were extracted from approximately 5–10 million cells using a modified acid extraction protocol. Cell pellets were first resuspended in Sucrose Buffer containing 0.3 M sucrose, 15 mM NaCl, 10 mM HEPES pH 7.9, 2 mM EDTA, 0.5% NP-40, and protease inhibitors. Nuclei were pelleted by centrifugation and washed twice in High Salt Buffer composed of 0.35 M KCl, 10 mM Tris-HCl pH 7.2, and 5 mM MgCl₂. The first high-salt wash was carried out with intermittent vortexing over 20 minutes to enhance chromatin solubilization. Following centrifugation, the nuclear pellet was resuspended in 0.2 N sulfuric acid (H₂SO₄), homogenized, and incubated overnight at 4 °C to release acid-soluble histones. The clarified supernatant was collected by centrifugation and subjected to precipitation with 10 volumes of 100% ice-cold ethanol.

Samples were incubated overnight at −20 °C, then pelleted by centrifugation, washed twice with 70% ethanol, air-dried, and resuspended in a minimal volume of nuclease-free water.

### HPLC Analysis of Histone proteins

Prior to chromatographic analysis, acid-extracted histones were filtered through a 0.2 µm Millex syringe filter directly into Waters screw neck polypropylene vials (12 × 32 mm, 250 µL capacity, Cat. No. WAT094172) to eliminate particulates and prevent bubble formation. Reverse-phase HPLC was performed using a Vydac 218TP C18 column (4.6 × 250 mm, 5 µm particle size, 300 Å pore size) on a Waters 2695 Separations Module. Samples were injected at a volume of 95 µL and separated using a linear acetonitrile gradient in 0.1% trifluoroacetic acid (TFA) over 93 minutes at a flow rate of 1 mL/min. The effluent was monitored at 214 nm using a Waters 996 Photodiode Array Detector.

For human histone samples, two methanol washes were applied to the column prior to injection to ensure baseline stability and reproducibility. Peak detection and quantification were performed using Waters Empower Pro software (version 2), and histone H1 peak areas were normalized to H2B peaks to account for sample input variability. Peak identification and quantification were carried out using Waters Empower Pro software (version 2). The relative abundance of histone H1 variants was determined by normalizing peak areas to H2B as an internal control for loading and recovery efficiency.

## RESULTS

### Histone H1.5 directly interacts with CENP-A mononucleosomes *in vitro*

Utilizing both FRET and FLIM methods, linker histone H1.0 and H1.2 closely associates with centromeric proteins CENP-A, CENP-B, and CENP-C in human cells (45). Linker histone H1.5 was reported to associate with centromeric chromatin arrays containing CENP-A -albeit at a lower affinity than H3-but poorly associates with GFP- or HA-tagged CENP-A *in vivo* (52). Recently, it was reported that histone H1.0 was able to interact with both H3 and CENP-A nucleosomes *in vitro* (48). Prior to that, isolation of CENP-A with various epitope tags yielded similar results of being unable to co-immunoprecipitate with linker histone H1s (44), consistent with recent findings that tagged CENP-A does not yield the same interactions as untagged native endogenous CENP-A *in vivo* (52).

To test whether native untagged CENP-A can form a complex with linker histones *in vitro*, we developed a method that allows for simultaneous detection of complexes containing both nucleic acid and protein. We reconstituted nucleosomes using salt dialysis of histone proteins (53) in the presence of high molecular weight (50,000–100,000) poly-L-glutamic acid (PGA) (as previously described in (54)) and 200 bp fragment containing the 601-positioning sequence with 5’ biotinylated ends (Fig. 1A-B). To confirm the presence of CENP-A-H1 mononucleosome complexes, reconstituted samples were electrophoresed to verify gel shift (EMSA), followed by native EMSA in-gel Western PAGE (NEW-PAGE) using antibodies against the histones and streptavidin-bound dyes targeting the biotinylated 200 bp PCR fragment (Fig. 1C). The unbound 200 bp PCR fragment (grey arrow) is shifted when nucleosomes are formed (red arrow) (Fig. 1C).

**Figure 1.**
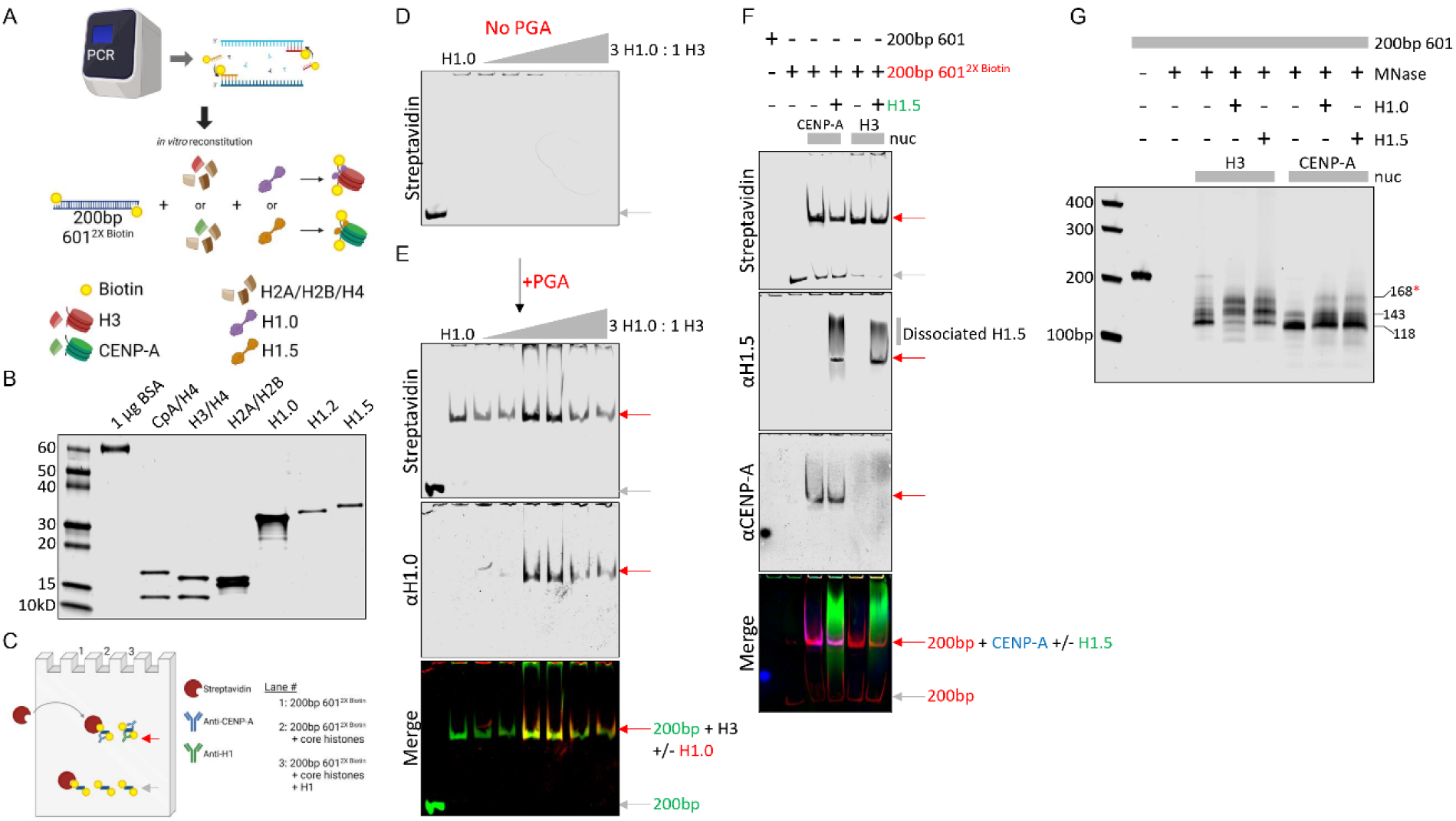
Histone H1.5 directly interacts with CENP-A nucleosomes *in vitro*. (A) Depiction of 200 bp PCR amplified 601 positioning sequence with 5’ biotinylated ends used in salt dialysis of H3 or CENP-A reconstitutions. (B) Some of the core and linker histones used in this study on a 4-20% SDS-PAGE gel. (C) Native EMSA in-gel Western PAGE (NEW-PAGE) protocol developed for this study to simultaneously detect biotinylated DNA (with streptavidin dye) and histones (with antibodies). (D) NEW-PAGE of H3 reconstitutions performed without PGA added in the presence of increasing H1.0 (0-to 3-fold excess over H3). (E) NEW-PAGE of H3 reconstitutions performed with PGA in the presence of increasing H1.0. (F) Reconstitutions of H3 or CENP-A nucleosomes with biotinylated 200 bp 601 positioning fragments in the absence or presence of H1.5, followed by NEW-PAGE co-immunofluorescence detection of DNA (streptavidin), H1.5, and CENP-A. (G) Reconstitutions of H3 or CENP-A nucleosomes with 200 bp 601 positioning sequence without H1.0 or H1.5, followed by MNase digestion to determine protection. Grey arrow = unshifted 200bp DNA fragment. Red arrow = shifted 200 bp fragment due to reconstituted nucleosome. *Red asterisk = additional MNase protected band when H1.0/H1.5 is added to CENP-A nucleosome.

In the absence of PGA, we observe large aggregates that were unable to migrate into the native PAGE (Fig. 1D). However, in the presence of PGA, histone H1.0, and H3 nucleosomes, we saw co-immunofluorescence of the shifted DNA fragment with both histones H1.0 and H3 (Fig. 1E).

Unlike H3, we observed no H1.0 interactions with CENP-A nucleosomes *in vitro* when using the 200 bp fragment with the 5’ biotinylated ends (Fig. S2A). However, when using the non-biotinylated ends, the H1.0-CENP-A nucleosome interactions were restored (Fig. S2B), suggesting the biotinylated ends interfere with H1.0 binding to CENP-A nucleosomes but not to H3.

Next, we reconstituted H3 or CENP-A nucleosomes in the absence or presence of linker histone H1.5 and observed that H1.5 forms a stable complex with the 200 bp biotinylated 601 fragment and CENP-A (Fig. 1F). Similarly, we see both CENP-A-H1.2 and CENP-A-H1.5 complexes on NEW-PAGE throughout the H1 titration series, suggesting both H1.2 and H1.5 can form a stable complex with CENP-A nucleosomes (Fig. S2C-D).

An alternative approach to multicolor in-gel co-detection of histones and DNA is to perform MNase digestion on reconstituted nucleosomes and examine the DNA protection pattern in the absence or presence of histone H1s. We observed an additional band above CENP-A control in the presence of H1.0 or H1.5, suggesting both linker histones are capable of protecting a larger DNA fragment when bound to CENP-A nucleosomes (Fig. 1G). Similarly, when we added linker histone H1.2 to CENP-A reconstitutions, we also observed an additional protected band post-MNase digestion, albeit a slightly shorter fragment (blue arrow, Fig. S2E) than H1.0 and H1.5 (red arrow, Fig. S2E), suggesting different linker histones will interact with CENP-A nucleosomes differently.

### *In vitro*, H1.5 associates with CENP-A and H3 mononucleosomes with distinct characteristics

Next, we wondered what the topographical consequences were of H1.5 binding to *in vitro* reconstituted CENP-A and H3 mononucleosomes. We designed a 324-bp DNA fragment with the nucleosome-positioning 601 DNA sequence at its center and two flanking sequences of differentiating lengths (68bp and 109bp, respectively) (Fig. 2A, S3A-B). We added H1.5 to half the sample at a ratio of one H1.5 molecule per one reconstituted mononucleosomes and incubated at room temperature for one hour. Next, we imaged the samples by AFM using tapping mode, allowing us to obtain topographical dimension of mononucleosome (Fig. 2B). *In vitro* reconstituted CENP-A and H3 nucleosomes have similar dimensions, based on their height measurements (2.35 ± 0.01 nm vs 2.38 ± 0.02 nm, respectively; Table 1, Fig. 2C) and Feret’s diameter (12.0 ± 0.1 nm vs 11.4 ± 0.2 nm, respectively; Table 1, Fig. 2D). When we added H1.5, we observed no change in height for H3 mononucleosomes (2.37 ± 0.02 nm; Table 1, Fig. 2C) but we observed a larger Feret’s diameter (11.4 ± 0.2 nm vs 15.4 ± 0.2 nm, respectively; Table 1, Fig. 2D), shorter DNA entry and exit strands (Fig. 2E), and tighter angles of the entry and exit strands (Fig. 2F). These changes in topographical features of H3 mononucleosomes upon H1.5 binding might indicate that H1.5 binds at the nucleosome dyad (39–42). In contrast, the Feret’s diameter for CENP-A mononucleosomes upon H1.5 binding did not change (11.7 ± 0.2 nm; Table 1, Fig. 2D), whereas CENP-A mononucleosomes bound to H1.5 were taller than CENP-A mononucleosomes alone (2.35 ± 0.01 nm vs 2.42 ± 0.02 nm, respectively; Table 1, Fig. 2C).

**Figure 2.**
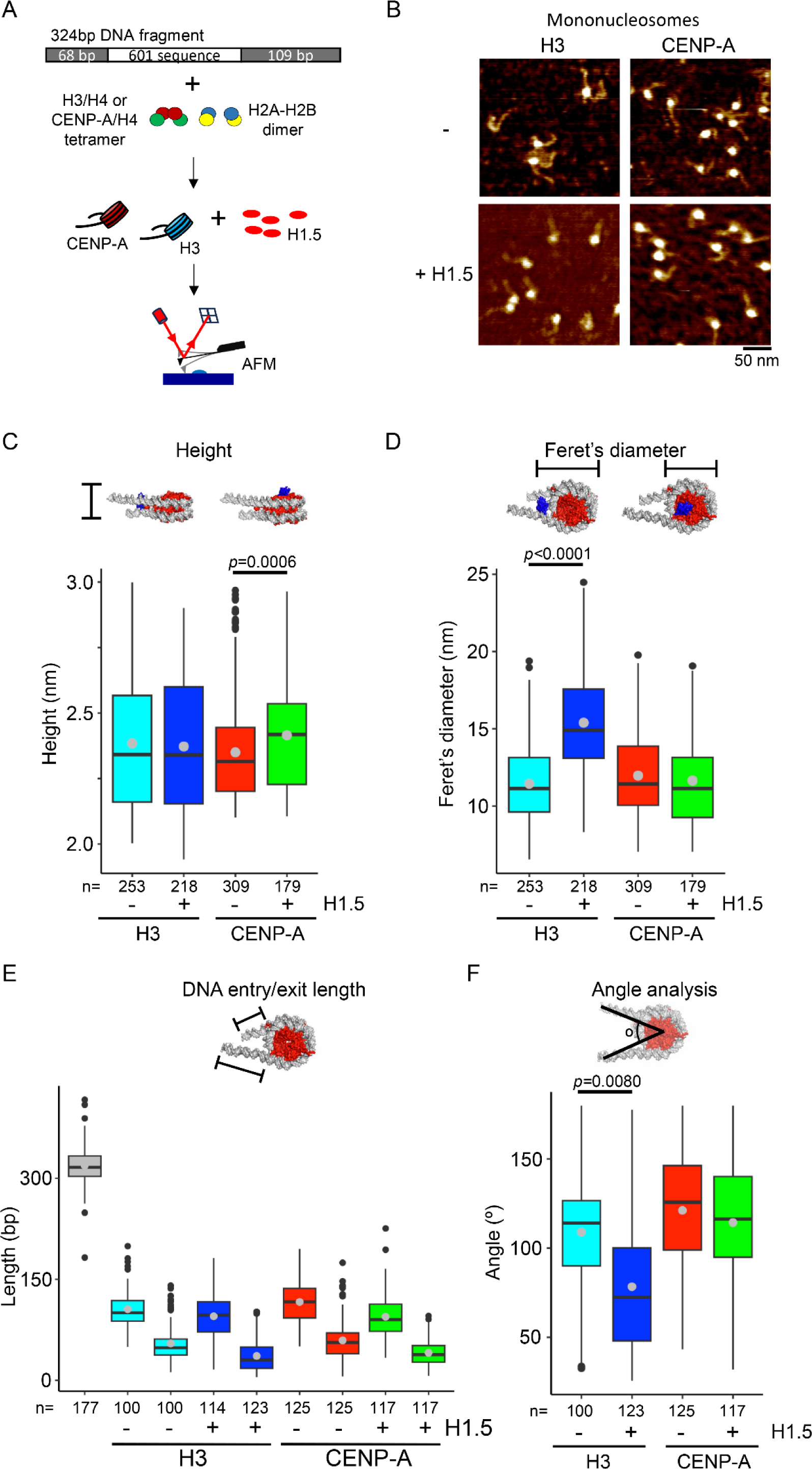
H1.5 might bind CENP-A mononucleosomes in a non-canonical manner. (A) Schematic representation of *in vitro* reconstitution of mononucleosomes followed by AFM imaging using tapping mode. (B) Representative AFM images of H3 and CENP-A mononucleosomes with and without H1.5. (C) Boxplot of nucleosome height measurements of H3 and CENP-A mononucleosomes with and without H1.5. (D) Boxplot of the Feret’s diameter measurements of H3 and CENP-A mononucleosomes with and without H1.5. (E) Boxplot of length of DNA fragment, as well as the long and short arm of the entry and exit DNA strands from the nucleosomes. (F) Boxplot of the angle between the long and short DNA arms of H3 and CENP-A mononucleosomes with and without H1.5.

**Table 1.** Quantification of recombinant nucleosome dimensions with H1.5.

| Nucleosome | n | Nuc height (nm) | Feret (nm) | n | DNA length (bp) |  | Angle |
| --- | --- | --- | --- | --- | --- | --- | --- |
| DNA fragment |  |  |  | 178 | 320±38 |  |  |
|  |  |  |  |  | Long | Short |  |
| CENP-A | 309 | 2.35±0.01 | 12.0±0.1 | 125 | 118±35 | 60±32 | 121±65 |
| CENP-A + H1.5 | 179 | 2.42±0.02 | 11.7±0.2 | 117 | 95±31 | 41±20 | 114±36 |
| H3 | 253 | 2.38±0.02 | 11.4±0.2 | 100 | 107±28 | 55±28 | 109±31 |
| H3 + H1.5 | 218 | 2.37±0.02 | 15.4±0.2 | 123 | 107±34 | 44±25 | 78±34 |

**Table 2:**
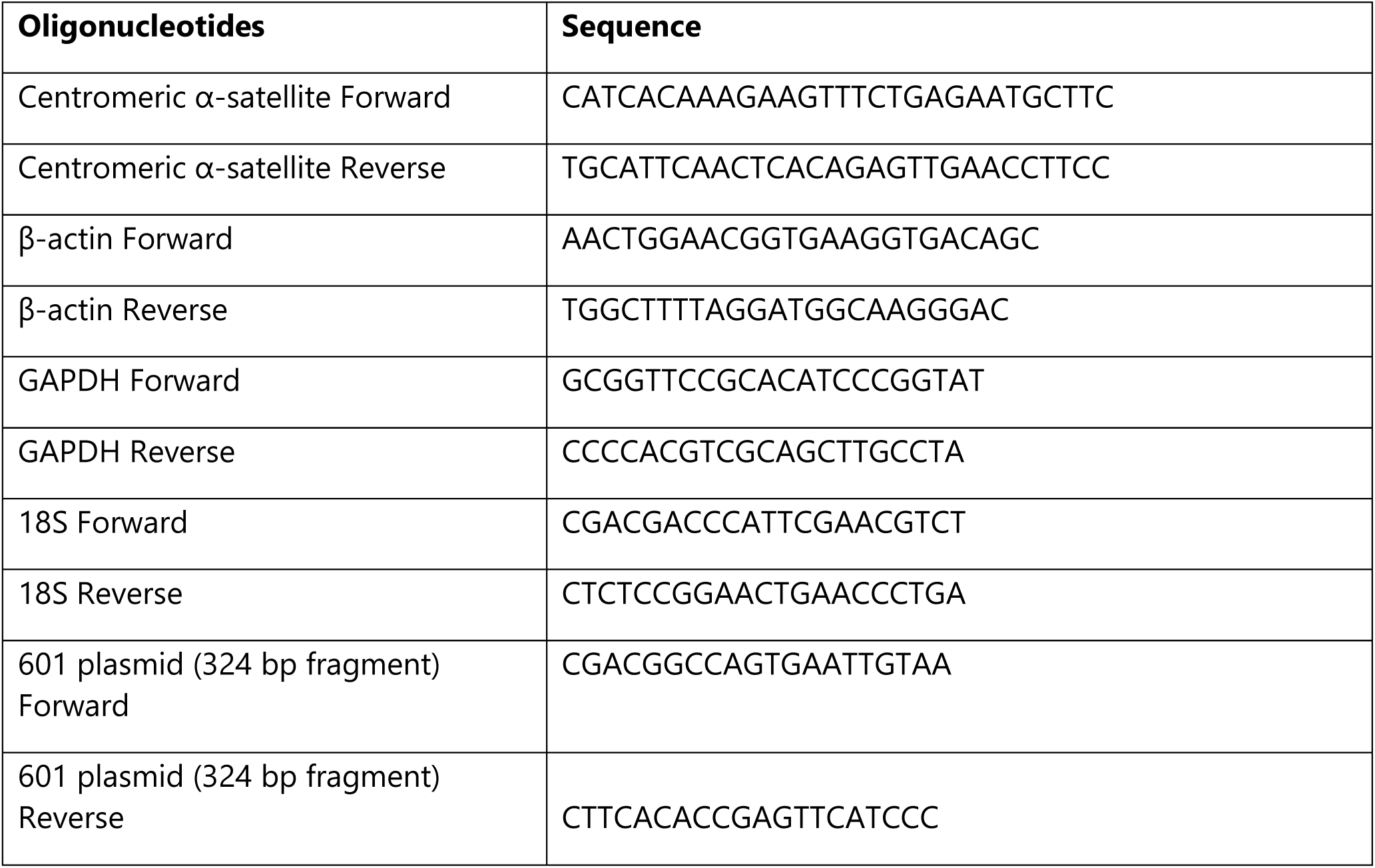
List of primer sequences used in this study.

Furthermore, the lengths of the entry and exit DNA strands did not alter when CENP-A was bound to H1.5 (Fig. 2E) or the angle between the entry and exit DNA strand (Fig. 2F). The observations of H1.5 increasing the height of CENP-A mononucleosomes, but not it’s Feret’s diameter, or the entry and exit DNA strand metrics, might imply that H1.5 does not sit near or at the dyad but instead that H1.5 binds CENP-A mononucleosomes independently from the dyad.

### Histone H1.5 localizes to centromeres *in vivo*

Previous reports (44, 45) and our *in vitro* findings (Fig. 1, 2) suggest a potential role for H1 at the centromere. Thus, we next sought to develop an *in vivo* test of this hypothesis.

First, to determine whether H1.5 localizes to centromeres, we performed immunostaining on normal diploid astrocytes using a custom affinity-purified antibody specific for H1.5. The specificity of the H1.5 antibody was confirmed using recombinant H1.5 versus other recombinant H1s (Fig. S4). We then used this validated anti-H1.5 antibody for immunofluorescence to detect H1.5 localization in cells, and co-stained for the centromeric marker Centromere Protein C (CENP-C). CENP-C is a well-established and conserved inner kinetochore protein that directly binds to CENP-A–containing nucleosomes and serves as a core component of the constitutive centromere-associated network (CCAN). Because of its stable association with functional centromeres throughout the cell cycle and its direct interaction with CENP-A, CENP-C is widely used as a reliable marker for centromere identity and localization (39, 41, 55). Our results showed that 65% complete co-localization and 28% partial co-localization of H1.5 and CENP-C foci (Fig. 3A,B). These data provide the first *in vivo* evidence that H1.5 localizes to centromeres.

**Figure 3.**
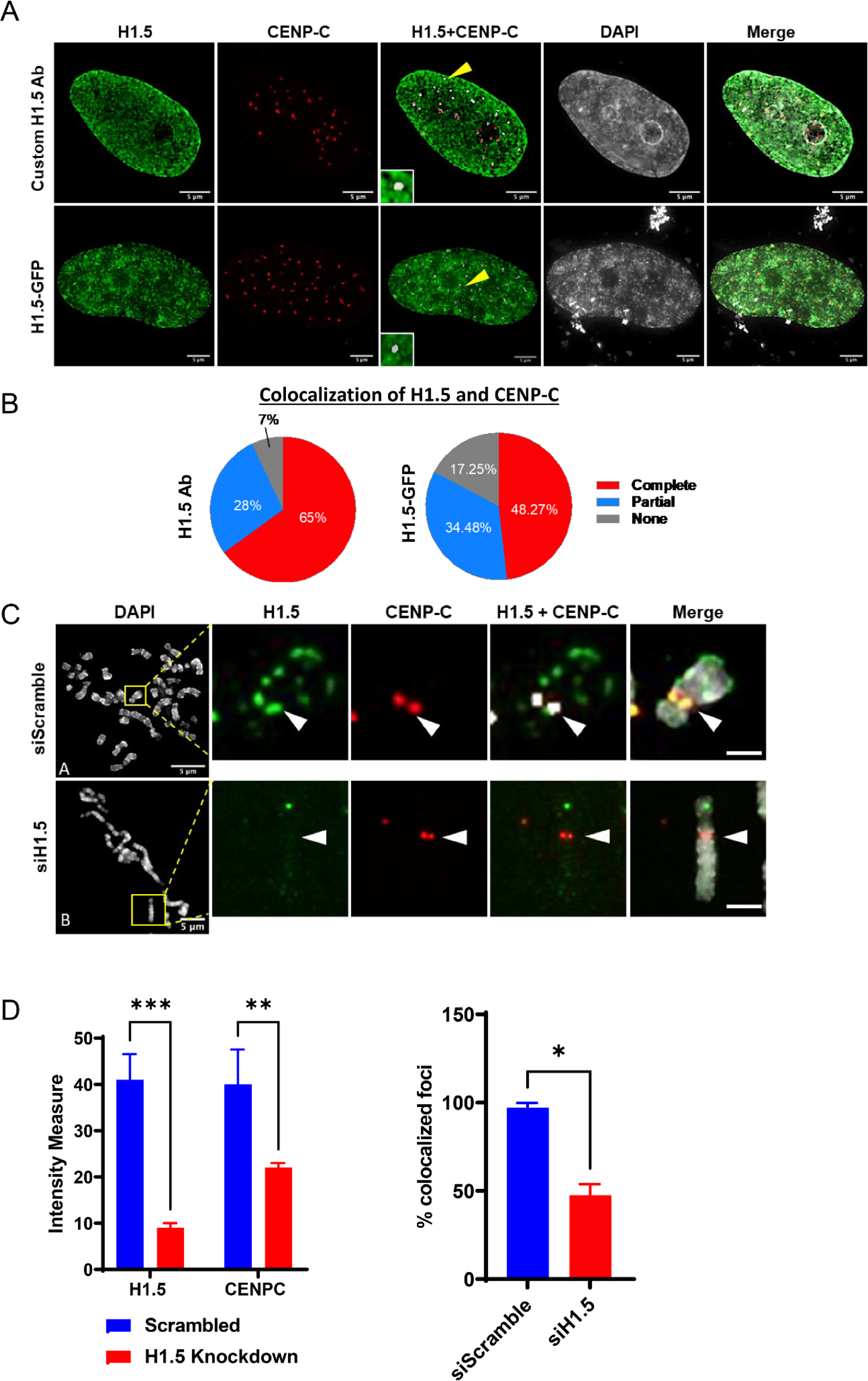
H1.5 co-localizes with the centromere marker CENP-C in interphase and metaphase cells. (A) Immunofluorescence images of SVGp12 interphase nuclei show colocalization between H1.5 (green) and CENP-C (red), visualized either using a custom H1.5 antibody (top row) or an H1.5-GFP fusion protein (bottom row). Overlap of the two signals is evident in the merged channels and enlarged insets. (B) Quantification of H1.5 and CENP-C colocalization in interphase nuclei using ImageJ’s Colocalization plugin. Pie charts represent the proportion of cells showing complete, partial, or no colocalization in cells stained with the antibody or expressing H1.5-GFP. (C) Immunofluorescence on metaphase chromosome spreads from siScramble- and siH1.5-treated cells reveals disrupted colocalization of H1.5 and CENP-C upon H1.5 depletion. Insets highlight representative centromeres where colocalization is either preserved (scramble) or lost (siH1.5). (D) Quantification of fluorescence intensity for H1.5 and CENP-C at centromeres in metaphase spreads (left panel), and percentage of centromeres showing co-localized signals (right panel). A significant decrease in both intensity and colocalization is observed following H1.5 knockdown. Data represent mean ± s.e.m; *p < 0.05, **p < 0.01, ***p < 0.001. Scale bars: 5 μm.

To further validate these findings, we utilized an antibody-independent approach by transfecting the astrocytes with H1.5-GFP. We found 48% complete co-localization and 34% partial co-localization of H1.5-GFP and CENP-C foci (Fig. 3A,B). Therefore, we find that the co- localization of H1.5 and CENP-C using the custom H1.5 antibody or H1.5-GFP transfected astrocytes to be comparable in interphase cells.

Next, we tested whether H1.5 also localized to the centromere in mitosis. To confirm that mitotic staining with our custom H1.5 antibody is specific, we knocked down H1.5 by siRNA. HPLC and RNA-seq analyses efficient knockdown of H1.5 mRNA, while other somatic H1s remain unchanged (Fig. S5). These data confirm that H1.5 is specifically depleted in response to siRNA treatment without affecting the overall stoichiometry of other H1 subtypes. After preparing metaphase spreads, we performed co-immunostaining using antibodies against H1.5 and CENP-C. The co-localization of H1.5 and CENP-C in the siScramble metaphase spreads provides further evidence that H1.5 is localized to centromeres (Fig. 3C-top panel), and that the co-localization of H1.5 with CENP-C (Fig. 3C-bottom panel), is diminished by half when H1.5 is knocked-down (Fig. 3D). Taken together, these observations support the hypothesis that histone H1.5 localized to centromeres in both interphase and mitosis.

### Histone H1.5 associates with CENP-A mononucleosomes *in vivo*

We previously established that purified CENP-A-containing chromatin arrays can associate with histone H1.5 (52). Because CENP-A and H3 domains alternate at the centromeres (56), this observation does not exclude the possibility that linker histones associate with neighbouring H3 nucleosomes and not directly with CENP-A at the centromere. To exclude these long range, indirect H1.5 interactions, we extracted MNase digested HeLa nuclei enriched for mononucleosomes (∼147bp), followed by native (unfixed) CENP-A ChIP to test whether H1.5 co-immunoprecipitated with CENP-A (Fig. 4A, S6A). H1.5 co-immunoprecipitates with CENP-A ChIP, indicating histone H1.5 directly associates with CENP-A nucleosomes *in vivo* (Fig. 4B). To test whether H1.5’s interaction with CENP-A nucleosomes depend on its C-terminal domain, we expressed an HA-tagged H1.5 construct lacking the C-terminal domain (H1.5^ΔCTD^ ^157–227^) in HeLa cells. Co-immunoprecipitation followed by Western blot analysis revealed that even in the absence of the CTD, H1.5^ΔCTD^ ^157–227^ retains its ability to associate with CENP-A (Fig. S6B-C). These findings confirm that the interaction between H1.5 and CENP-A is maintained independently of the C-terminal domain and underscore a direct association between the two proteins *in vivo*.

**Figure 4.**
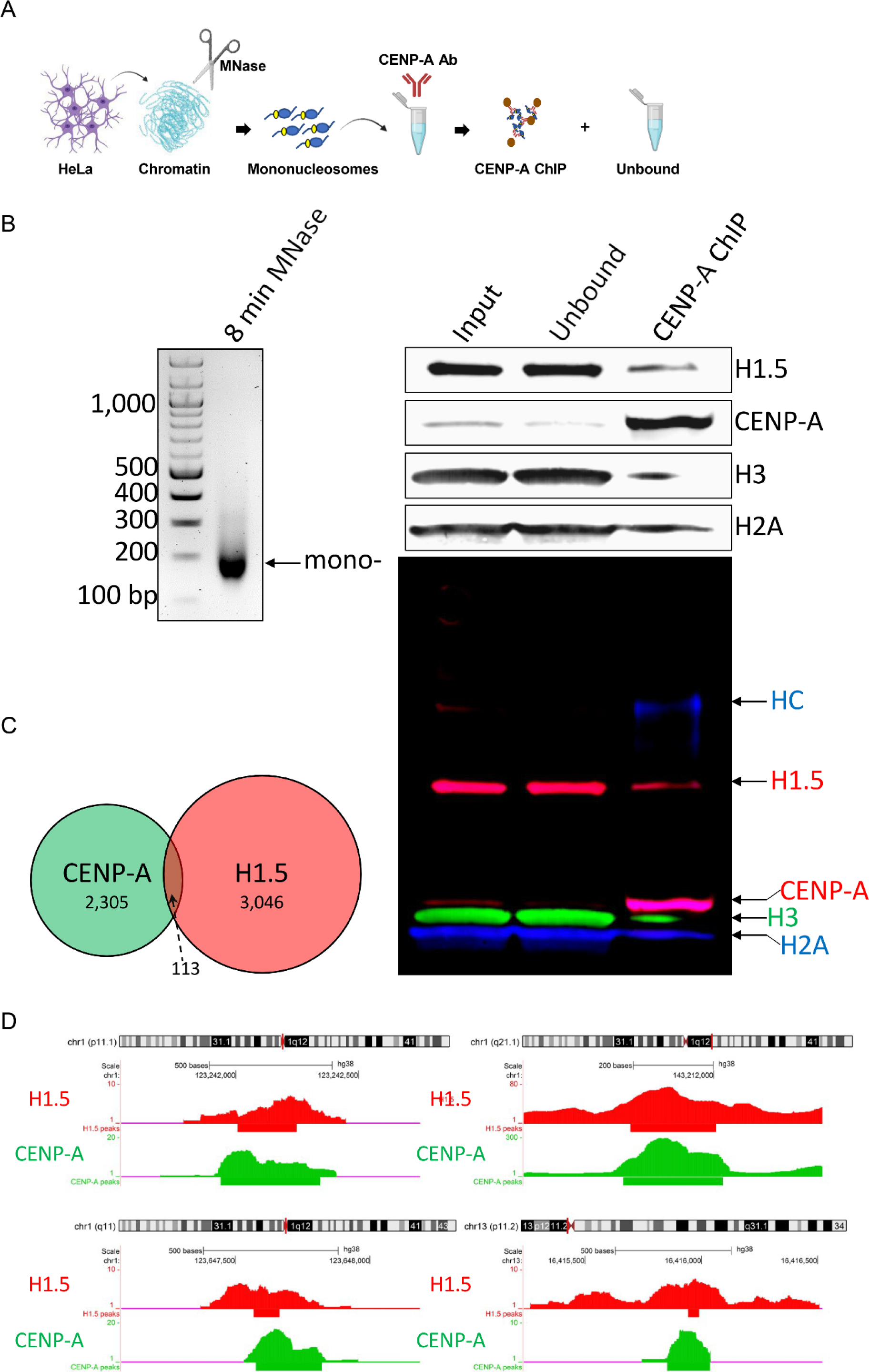
H1.5 interacts with CENP-A mononucleosomes, *in vivo*. (A) Schematic depicting MNase digestion of HeLa cells followed by CENP-A ChIP. (B) MNase ladder depicting digestion of chromatin to mononucleosomes. ChIP followed by Westerns against H1.5, CENP-A, H3, and H2A histones. 2.5% Input/Unbound and 100% CENP-A ChIP samples loaded. Unbound = supernatant not bound to CENP-A antibody/beads complex. HC=heavy chain of anti-CENP-A ChIP antibody. (C) Venn diagram comparing total shared peaks between native CENP-A and H1.5 histones. (D) Genome browser snapshots of overlapping native CENP-A and H1.5 peaks.

Another approach to investigate whether histone H1.5 and CENP-A occupy common genomic regions, we compared two publicly available ChIP-seq datasets from HeLa and T47D cancer cell lines, one generated in our lab for native CENP-A (52), and a previously published dataset for H1.5 (57), respectively. This comparative analysis revealed 113 genomic sites that are co-occupied by both CENP-A and H1.5, suggesting the possibility of their physical association at a subset of chromatin regions (Fig. 4C, D).

### H1.5 knock-down represses α-satellite transcription and CENP-A loading

H1s are known for their role in compacting chromatin and repressing transcription. Therefore, the presence of linker histone H1.5 at centromeres (Fig. 3-4) led us to speculate that H1.5 may facilitate centromeric transcription. The centromere is enriched with α-satellite DNA and its transcription was previously shown to be essential for CENP-A loading (58), and that its loss could result in poor kinetochore formation, resulting in mitotic defects (58, 59). To assess whether H1.5 depletion could impact α-satellite transcription universally across multiple cell lines, we performed qRT-PCR 3-days post-siRNA knock-down on SW480: metastatic colorectal cancer cells, HeLa: cervical cancer cell line, SVG: normal astrocytes, and BJ: normal fibroblasts. α-Satellite transcript levels were lowered across all cell lines upon SMARTpool siH1.5 knock-down, but most prominently among cancer cell lines SW480 and HeLa cells (Fig. 5A). A possible explanation for minimal α-satellite reduction in BJ cells post-siH1.5 knockdown might be that α-satellite transcript levels have a low base-line level (Fig. 5A) or that H1.5 may not serve as the major H1 in that cell line.

**Figure 5.**
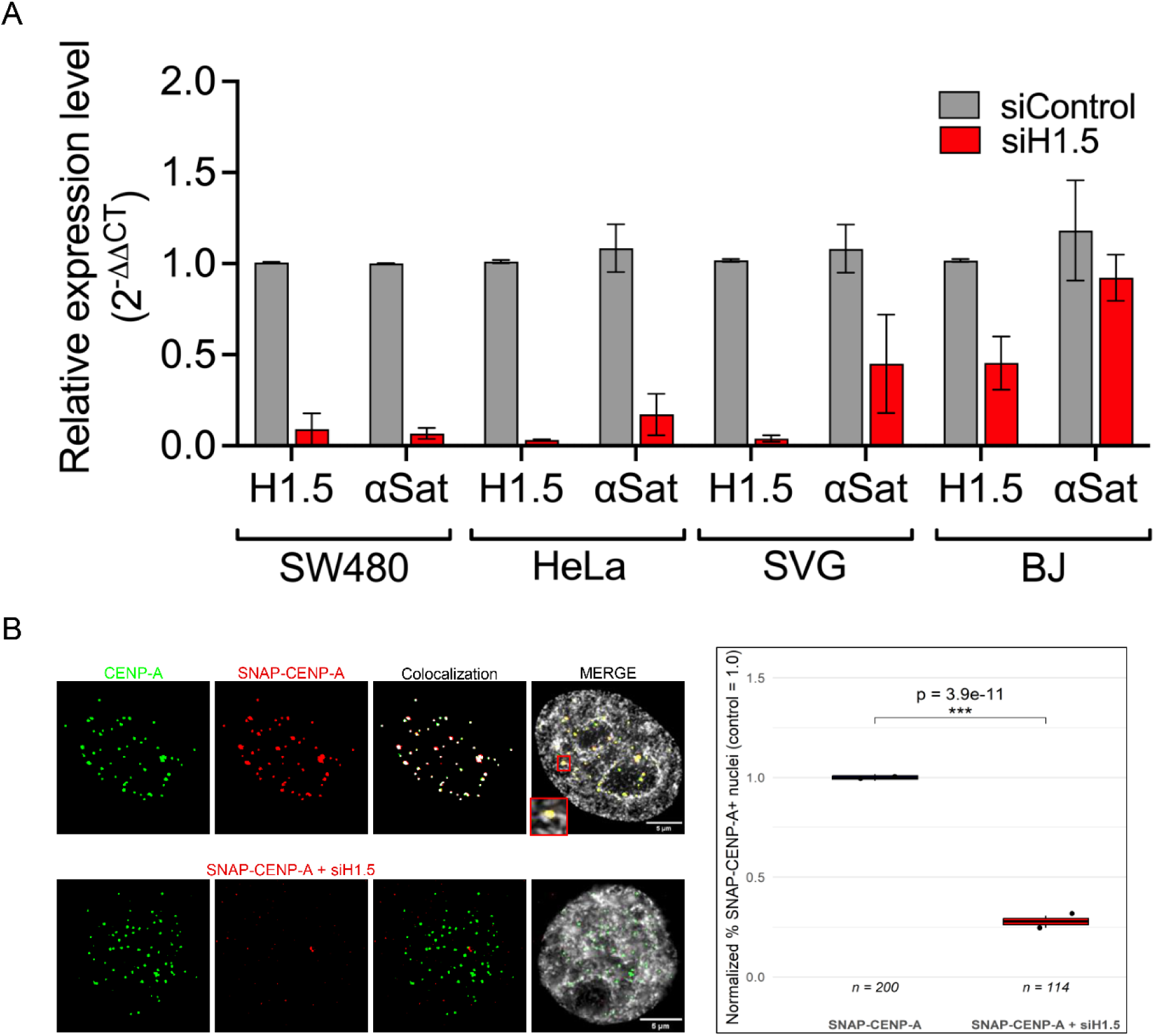
H1.5 knockdown reduces α-satellite transcription and impairs new CENP-A loading at centromeres. (A) Quantitative RT-PCR analysis of H1.5 and α-satellite (αSat) RNA levels across four human cell lines (SW480, HeLa, SVG, and BJ) treated with siScramble or siH1.5. H1.5 knockdown results in significant reduction of αSat transcription in SW480 (p < 0.01), HeLa (p < 0.01), and SVG (p < 0.001) cells, but not in BJ cells. Data represent mean ± SEM from at least two biological replicates. (B) Centromeric CENP-A loading assay using SNAP-tagged CENP-A. HeLa SNAP-CENP-A cells were synchronized to early G1 phase, labelled with TMR-Star to detect newly deposited CENP-A (red), and co-stained for endogenous CENP-A (green). Top row shows control cells with robust SNAP-CENP-A and native CENP-A colocalization. Bottom row shows reduced colocalization following siH1.5 treatment. SNAP-CENP-A was expressed in cells treated with control or siH1.5, and the percentage of nuclei showing >25% centromeric colocalization was measured. Values were normalized to the mean incorporation in the control condition (SNAP-CENP-A = 1.0). Each dot represents one biological replicate (n = 2 per condition), derived by splitting the total number of analyzed nuclei into two matched subgroups. Boxplots represent the distribution of normalized % SNAP-CENP-A+ nuclei across replicates. A significant reduction in incorporation was observed upon H1.5 knockdown (Fisher’s exact test, ***p = 3.9 × 10⁻¹¹).

To further evaluate the transcriptional specificity of H1.5 knockdown, we performed RNA-seq data from H1.5 knockdown astrocytes. While α-satellite transcription was significantly reduced, which was validated by qRT-PCR, global levels of noncoding RNAs (ncRNA) was upregulated (Fig. S7A-C). Interestingly, several major classes of repetitive element transcripts including LINEs, SINEs, and telomeric RNAs, remained unchanged (Fig. S7C). These results indicating that H1.5 predominantly represses ncRNA transcription, but with a notable exception of centromeric α-satellite transcription. To assess whether this effect was unique to H1.5, we also analysed RNA-seq data from siH1.0-treated cells. In contrast to H1.5 depletion, H1.0 knockdown did not affect α-satellite RNA levels but instead resulted in a modest decrease in global noncoding RNAs, with repetitive RNA categories such as LINEs, SINEs, and telomeric RNAs remaining unchanged (Fig. S7A, C). Furthermore, we observed that H1.5 depletion also reduced CENP-A expression across several cell types (Fig. S7D), which could contribute to insufficient or weakened kinetochores. BJ was the only cell line to overexpress CENP-A upon siH1.5 knock-down, suggesting H1.5 may regulate CENP-A gene expression differently in cells with low baseline levels of CENP-A (60).

If both α-satellite and CENP-A transcripts are reduced, the likely result is insufficient loading of new CENP-A histones during early G1. To assess whether new SNAP-CENP-A is loaded after early G1-phase, we transfected HeLa cells with SNAP-CENP-A and treated with either siScrambled control or siH1.5 knockdown, and followed by a double thymidine and TMR block/release. TMR staining against the SNAP-tag allowed us to visualize newly loaded SNAP-CENP-A at centromeres when co-stained with native CENP-A. We observed a 4-fold reduction in new SNAP-CENP-A being loaded in the H1.5 knockdown cells compared to the control cells (Fig. 5B).

To further investigate the transcriptional consequences of H1.5 depletion, we performed RNA-sequencing comparing either siH1.0 or siH1.5 to siScramble conditions in astrocyte cells. The volcano plot for H1.5 knockdown (Fig. S1D) revealed a distinct transcriptional profile with a substantial number of differentially expressed genes. Subsequent pathway and gene-level analyses showed strong downregulation of centromere- and mitosis-associated genes such as CENP-A, NUSAP1, and BUB3, alongside selective upregulation of mitotic regulators including PLK1, AURKA, and TPX2 (Fig. S8B-D). In contrast, the volcano plot for H1.0 knockdown (Fig. S8A) displayed a broader and more diffuse transcriptional profile with 1,033 upregulated and 705 downregulated genes (p-value <0.05, Table S1). Several mitotic regulators were affected in both knockdowns, but with different patterns: H1.0 knockdown broadly upregulated genes associated with chromosome segregation (Fig. S8C, D), while H1.5 knockdown showed downregulation of many centromeric components and upregulation of genes linked to mitotic progression (Fig. S8B, D).

Gene ontology analysis revealed that in both H1.0 and H1.5 knockdown the most enriched pathway was ‘cytoplasmic translation’. In addition, in the H1.0 knockdown was various ‘chromosome segregation’ pathways were upregulated, while H1.5 knockdown showed downregulation of categories such as ‘negative regulation of cell cycle’ (Fig. S8C). These enrichments underscore the distinct transcriptional consequences of H1.0 versus H1.5 depletion during mitotic progression. We then performed a targeted analysis focusing on mitotic gene sets from the Reactome “M Phase” and “Mitotic Metaphase and Anaphase” pathways in the Molecular Signatures Database (MSigDB) (60). This analysis showed that the mitotic kinases PLK1, AURKA, and AURKB were upregulated in both knockdowns, whereas centromeric genes including BUB3, NUSAP1, and CENP-B were downregulated only in H1.5-depleted cells, consistent with a centromeric role (Fig. S8B, D).

These observations complement our findings that H1.5 depletion reduces α-satellite transcription and impairs CENP-A loading, suggesting that H1.5 may contribute more broadly to the transcriptional coordination of centromeric and mitotic genes.

### H1.5 knock-down results in mitotic defects and altered CENP-A expression

Since our study demonstrates that histone H1.5 associates with CENP-A at the centromere and knocking down H1.5 resulted in reduced centromeric transcription and impaired loading of new CENP-A, we sought to determine whether depleting H1.5 *in vivo* would affect centromere function. To investigate this, we analyzed the impact of H1.5 knockdown on mitotic progression and mitotic defects.

Cells were arrested at G1/S by a double thymidine block and collected every 3 hours post-release over a 24-hour period. The samples were then analysed for DNA-content with flow cytometry (FACS) to track the cell cycle (Fig. S9). Both siScramble and siH1.5 treated cells reached G1 by 12-hour post-release. However, we observed a lag in the cells progressing between replication and G2/M, specifically between the 3- to 9-hour time point.

Next, we asked whether H1.5 depletion affected the ability of the cells to undergo mitosis. After releasing the double thymidine blocked for 9 hours, the cells were collected and subjected to co-IF staining (Fig. 6A) for CENP-C and α -tubulin (α-tub). The samples were quantitatively assessed for mitotic defects (Fig. 6B). 93% of the siScramble-treated/control cells display a typical metaphase pattern where we observe symmetrical alignment of centromeres at the metaphase plate, mitotic spindle fibers arranged uniformly (61). However, H1.5 knockdown revealed three distinct mitotic defects; (i) misaligned metaphase plates (ii) occurrence of multipolar spindles (iii) presence of lagging chromosomes (Fig. 6B). The most prominent phenotype (52%) was where the centromeres, marked by CENP-C, were found distributed along the spindle fibers (Fig. 6B).

**Figure 6.**
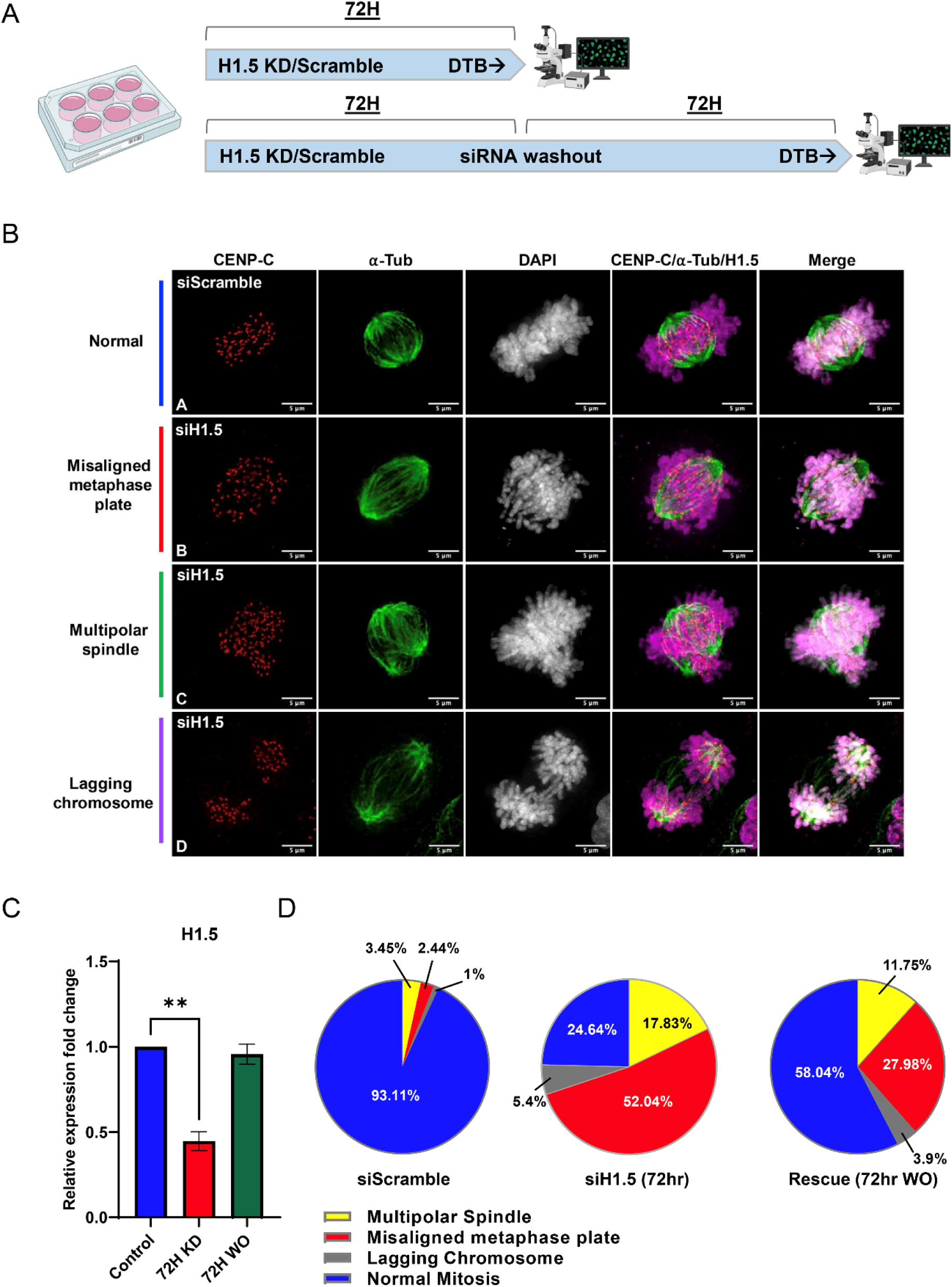
Loss of H1.5 leads to mitotic defects in normal human astrocyte cells (SVGp12). (A) Schematic of the siRNA knockdown and washout experimental design. SVGp12 cells were treated with either siScramble or siH1.5 for 72 hours, followed by a double thymidine block (DTB) to enrich for mitotic populations prior to fixation and immunofluorescence analysis. For the washout experiment, cells were cultured for an additional 72 hours in siRNA-free media before synchronization and fixation. (B) Representative immunofluorescence images of mitotic phenotypes observed in SVGp12 cells stained for centromeres (CENP-C, red), microtubules (α-tubulin, green), and DNA (DAPI, gray). siScramble-treated cells exhibit normal mitotic progression (Row A). In contrast, siH1.5-treated cells display a range of mitotic defects including misaligned metaphase plates (Row B), multipolar spindles (Row C), and lagging chromosomes (Row D). (C) RT-qPCR quantification of H1.5 mRNA levels confirms efficient knockdown by siH1.5 and partial restoration of expression following washout. Bars represent mean ± SEM from biological replicates. (D) Quantification of mitotic defects in >100 cells per condition. H1.5-depleted cells show significantly increased frequencies of multipolar spindles (p < 0.001), misaligned metaphase plates (p < 0.01), and lagging chromosomes (p < 0.01) compared to siScramble controls. Partial rescue of mitotic phenotypes is observed upon H1.5 re-expression following siRNA washout.

Finally, we asked whether restoring H1.5 levels could reverse these mitotic defects. To this end, we performed a rescue experiment where we washed out the siH1.5 following the 72hr knock-down and allowed the cells to grow in fresh culture media for an additional 72hrs to allow H1.5 mRNA levels to normalize (Fig. 6C), before cell synchronization and immunostaining.

Interestingly, all three of the observed defects were partially reversed where the most prominent rescue that of the misaligned chromosomes at the metaphase plate (28% vs 52%, respectively) (Fig. 6D). These observations suggest H1.5 plays a vital role in cell cycle progression through mitosis.

## DISCUSSION

Linker histone H1 variants in mammals play essential roles in shaping chromatin structure and regulating gene expression, often coordinated with developmental time. Indeed, more than 20 years ago, seminal work from the Skoultchi lab demonstrated that combinatorial deletion of H1 variants leads to variant-specific defects, and even lethality in mice (24, 25, 62). While traditionally regarded as global repressors of transcription, emerging evidence suggests that H1 subtypes may carry out distinct regulatory functions depending on their genomic localization such as transcriptional activation (63). Yet the contributions of these fascinating, highly positively charged proteins at specialized chromatin domains such as the centromeres remained a tantalizing challenge (64). In this study, we systematically studied this problem using interdisciplinary approaches, leading to the discovery of a distinct role for histone H1.5 at human centromeres in diploid glial cells.

Using IF (Fig. 3) and ChIP-seq (Fig. 4), we observed that H1.5 is enriched at centromeres. Targeted depletion of H1.5 leads to a marked loss of centromeric α-satellite transcription and a corresponding reduction in CENP-A expression on human centromeres. Disruption of H1.5 is functionally consequential, resulting in mitotic defects such as chromosome misalignment, lagging chromosomes, and spindle abnormalities. Further, two types of *in vitro* assays- an adaptation of the fluorescence-based EMSA assay first developed by the Luger lab (65), and single molecule imaging by AFM (53, 66) confirm the physical association of H1.5 with CENP-A mononucleosomes. Interestingly, when examining whether linker histone H1.0 and H1.2 also forms a complex with CENP-A, we observed H1.0 interaction only when utilizing a non-biotinylated 200 bp DNA fragment (Fig. S2A-B), suggesting the biotinylated 5’-end interferes with H1.0 binding. H1.2 also interacts with CENP-A but its MNase protected DNA signature differs from both H1.0- and H1.5-bound nucleosomes (Fig. S2E). The data indicates that linker histone H1s interact with CENP-A and form structures that are distinct from canonical histone H3 nucleosomes, which is further supported by the difference in height, Feret’s diameter, and angles observed between H3- and CENP-A-nucleosomes in the absence or presence of histone H1.5 (Fig. 2).

We next examined whether linker histones contributed to mitotic regulation by knocking down histone H1.0 and H1.5. The distinct distribution of mitotic defects observed upon H1.0 (Fig. S10) and H1.5 (Fig. 6) knockdown suggests that these two linker histone variants may regulate different aspects of mitotic chromosome architecture and spindle dynamics. The high frequency of misaligned metaphase plates in H1.5-depleted cells (52%) points to a potential role for H1.5 in ensuring proper kinetochore-microtubule attachment or in maintaining centromere architecture necessary for accurate chromosome alignment (Fig. 6). This is consistent with our finding that H1.5 directly associates with CENP-A-containing nucleosomes and is enriched at centromeres.

Conversely, the predominance of multipolar spindles upon H1.0 knockdown (41.6%) could indicate a broader role for H1.0 in centrosome organization, spindle pole integrity, or microtubule nucleation (Fig. S10). Since H1.0 has been shown to have a more global chromatin-binding profile, its depletion likely is complicated by widespread chromatin decompaction that disrupts overall nuclear organization, thereby, we speculate, indirectly affecting spindle assembly.

Together, these observations raise an interesting possibility that linker histone variants do not always act redundantly as first demonstrated by the seminal depletion studies in mice (24, 25, 67) but may contribute uniquely to chromosome segregation -H1.5 through centromeric chromatin regulation, and H1.0 potentially via more global chromatin or nuclear structural roles.

In addition to reducing CENP-A at centromeres, we also found that H1.5 knockdown resulted in transcriptional changes affecting several other centromeric components. RNA-seq analysis revealed a downregulation of some centromeric genes such as CENP-A, CENP-M, CENP-B, CENP-C, CENP-S, and CENP-F and upregulation of other centromeric genes such as CENP-N, CENP-O, and CENP-I (Fig. S8), which may reflect a compensatory response to centromere destabilization (68). Furthermore, mitotic checkpoint regulators PLK1, AURKA, and SPDL1 were significantly upregulated, suggesting activation of the spindle assembly checkpoint (69). Although global gene ontology analysis did not highlight enrichment in mitotic pathways when H1.5 was knocked down, a focused gene set approach using curated “M Phase” and “Mitotic Metaphase and Anaphase” signatures from the Molecular Signatures Database (MSigDB) (70) revealed consistent differential expression of mitotic and centromere-associated genes. These findings might contribute to the phenotypic consequences of H1.5 depletion, suggesting a role in the broader transcriptional coordination necessary for mitotic fidelity.

Our comparative RNA-seq analysis of H1.0 and H1.5 knockdown further supports this functional distinction. While H1.5 knockdown resulted in pronounced transcriptional alterations, including selective down- and upregulation of centromeric and kinetochore genes (Fig. S8), H1.0 knockdown affected a set of genes different from H1.5 knockdown, such as upregulation of genes involved in chromosome segregation including various mitotic kinases. This highlights the unique role of H1.5 in maintaining centromeric transcription and mitotic fidelity, distinguishing it from other somatic H1 variants.

Examining reconstituted recombinant CENP-A mononucleosomes *in vitro* suggests a novel binding mode for H1.5, where the globular domain binds centromeric nucleosome *independent of the dyad* (Fig. 7). H1.5 is typically thought to bind canonical H3 nucleosomes using both its globular domain (GD) and C-terminal domain (CTD), which together stabilize the protein at the nucleosomal dyad and contribute to chromatin compaction (12). This unusual observation points to exciting new –and potentially dynamic-binding modes for specific H1 variants. For example, based on our H1.5:CENP-A dyad-independent interpretation, dyad-proximal histone modifications might impact H1.5 binding (50, 71), which may impact access to internal nucleosome residues that are normally sterically hindered, in a manner reminiscent of recent outstanding work on HP1 “melting” the octameric core of the H3 nucleosome (72). In the context of this study, we speculate that a dyad-independent binding mode of H1.5 might promote RNAP

**Figure 7.**
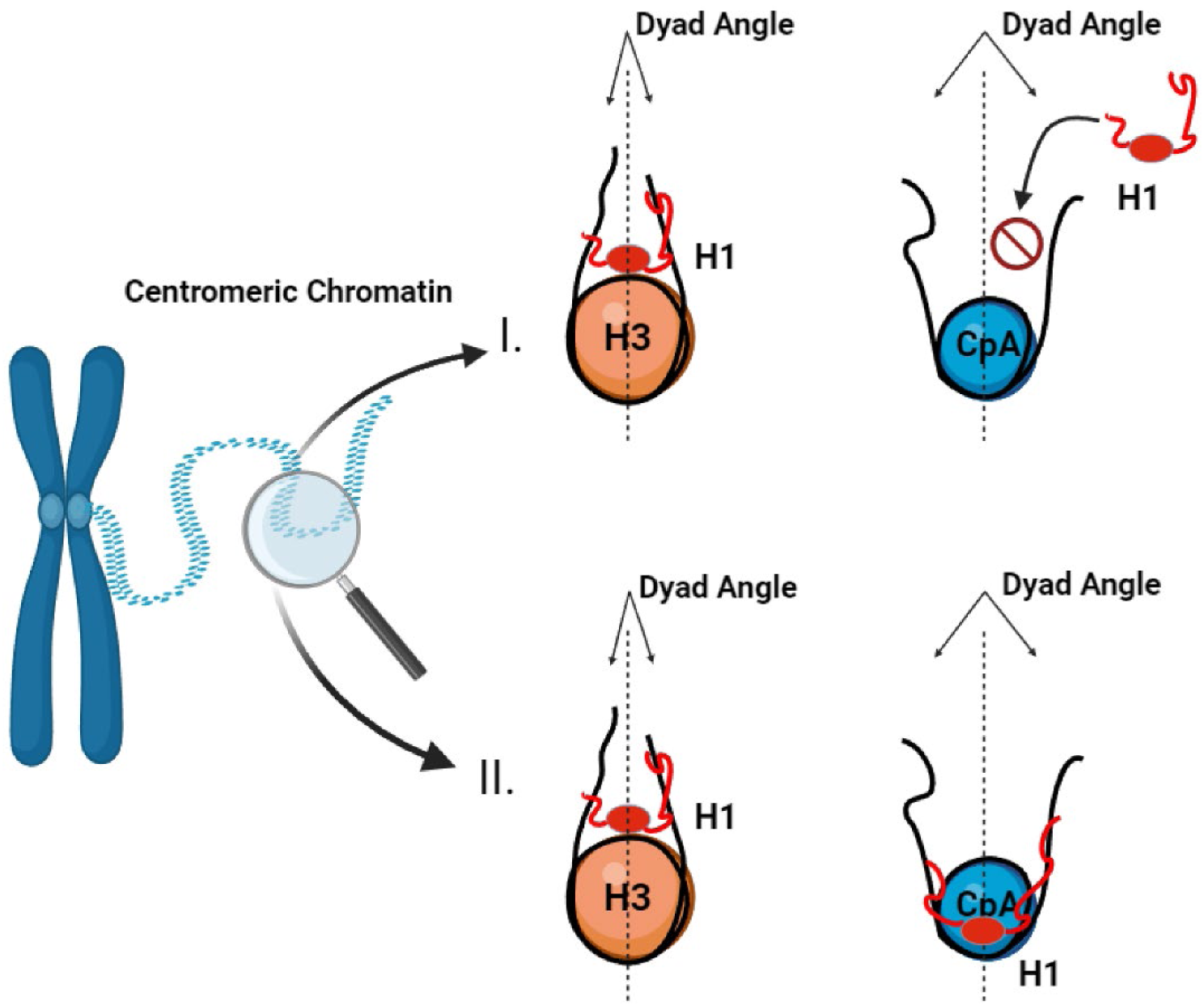
Proposed conceptual model for H1.5 interaction with centromeric chromatin and its role in mitotic fidelity. Schematic comparing canonical and proposed modes of H1.5 interaction with nucleosomes. (I) Canonical histone H1 is known to bind H3 nucleosomes at the nucleosomal dyad, but is precluded from CENP-A nucleosomes due to altered dyad geometry and increased DNA flexibility. (II) Based on our *in vitro* findings and supported by prior computational modelling, we propose that H1.5 may engage CENP-A nucleosomes through a non-canonical, dyad-independent mode of binding that permits its stable association with centromeric chromatin.

II mediated centromeric transcription, instead of the expected repressive effect typically associated with H1-nucleosome binding (see Graphical abstract). This hypothesis, we hope, can be tested in future structural studies. It has been previously reported that centromeric transcription is necessary for CENP-A loading (reviewed in (59)) Therefore, if this process is perturbed, we expect a reduction in newly loaded CENP-A. To test this, we transfected and synchronized SNAP-CENP-A cells to post-early G1, and co-stained with native CENP-A and TMR-star (against the SNAP-tag). Our results indicate a 4-fold reduction of newly loaded SNAP-CENP-A when H1.5 is knocked down compared to siScramble (Fig. 5B), indicating H1.5 plays a role in both centromeric transcription (Fig. 5A) and new CENP-A loading (Fig. 5B).

It is interesting to note that a prior study conducted using *Xenopus* oocytes where each somatic H1 was overexpressed, H1.5 was the only subtype that did not result in a saturable increase in nucleosome repeat length (NRL) (73). The change in NRL caused by the other somatic H1s was separately reported in a triple knockout mouse embryonic stem cell system where the loss of H1 was accompanied by a decrease in NRL (25). Therefore, given the unusual binding mode proposed in this study, we speculate that H1.5 may be hindered in its ability to alter nucleosome spacing due to its flexible interaction with chromatin, resulting in multiple conformations (Fig. 7).

To better understand the structural requirements for this interaction, we examined a C-terminal deletion mutant of H1.5^ΔCTD^ ^157–227^, which lacks the domain typically required for nucleosome binding. Surprisingly, H1.5^ΔCTD^ ^157–227^ still bound CENP-A nucleosomes, suggesting a non-canonical mode of engagement. Supporting this, AFM revealed that H1.5 binds CENP-A nucleosomes in a manner distinct from its dyad-centered interaction with H3 nucleosomes (Fig. 2). Although DNA wrapping can influence nucleosomal height as measured by AFM (74), we also measured the angle of the entry and exit DNA strand and found that only when H1.5 was bound to H3 nucleosomes the angle was altered (Fig. 2F). Taken together, these findings point to a unique, dyad-independent binding mode that may underlie H1.5’s specialized function at centromeres. Literature has been ambiguous about the possibility of H1 at the centromere, as several studies examining how H1s can directly bind CENP-A nucleosomes have been contradictory. The basis for this specific interest is that CENP-A wraps less DNA than a canonical H3 nucleosome (75). An early report using FLIM and FRET on HeLa cells detected the presence of some H1 subtypes, specifically H1.0 and H1.2, at centromeric chromatin with CENP-A, CENP-B, and CENP-C (45). However, cryo-EM and hydroxy-OH foot-printing showed a weak/no interaction between CENP-A and H1.5, and concluded that the increased flexibility of the linker DNA arms of CENP-A nucleosomes preclude the binding of H1 (14). A recent AFM study on H3 and CENP-A mononucleosomes showed that H1.0 can bind both nucleosomes, albeit in different modes. H1.0 binds H3 nucleosomes at the dyad, whereas H1.0 binds CENP-A by binding the entry and exit DNA strands (48). The disorganized nature of the NTD and CTD of H1, and H1 binding studies limited to the globular domain may have contributed to the assumption that only dyad-dependent binding was possible. With recent advances made in solving the dynamics of the H1 terminal domains (76), it is possible that further patterns may be observed. The H2A-H2B acidic patch is recognized by most chromatin-binding proteins (77, 78). Here, we postulate that this alternative H1 binding mode to the H2A acidic patch might be the preferred binding more for H1.5 to CENP-A nucleosomes. One potential consequence of this binding mode might be that H1.5 creates a steric hinderance for factors that bind to the H2A acidic patch.

It is established that H1:protein interactions regulate higher order structures, such as H1 recruiting Su(var)3-9 to establish HP1-type heterochromatin (27) and pioneer factor FOXA1 displacing H1 to open compacted chromatin to permit transcription (79, 80). There is growing evidence that H1s can also mediate chromatin-coupled protein-protein interactions (reviewed in (81)). Among the various heterochromatin domains, H1s also localize to the nucleolus (82, 83), where H1.0 can bind factors that regulate in rRNA transcription (84–86). It will therefore be of interest to study if H1s can recruit chromatin proteins to the centromere.

Finally, it is now accepted that H1s, like core histones, are subjected to post-translational modifications (PTMs). These PTMs can alter the charge of the amino acid residues, thus impacting H1 residency time on chromatin and its interaction with other proteins. For example, phosphorylation of H1.2 at serine 172 and H1.5 phosphorylated at serine 172 leads to their localization to sites of active DNA replication and transcription (87). H1.2 phosphorylated at serine 173 is associated with active transcription as it is enriched in interphase nucleoli and associated with transcribing rDNA, likely acting as a facilitator of RNAPI transcription (88). Histone H1.4 phosphorylated at serine 187 was also found to be associated with RNAPII mediated transcription in a hormone induced model system (63). Therefore, it is possible that other H1 variants with altered PTM status may display a binding pattern similar to H1.5 observed in this study.

CENP-A is an indispensable component for centromere specification, sufficient to seed and propagate a functional centromere/kinetochore (39, 89–92), and is epigenetically responsible for the formation of new centromeres at distal sites. Neocentromeres are functional centromeres that are formed at ectopic sites either due to disruption of the original centromere, or in cancer cells where CENP-A is overexpressed (39, 60, 93, 94). The discovery of these structures suggests that a specific DNA sequence, namely α-satellite sequence, is not required for centromere function. An exciting future avenue of research is to investigate the roles of H1s at neocentromeres, which also depend on noncoding transcription or distorted chromatin structures such as R-loops. Together, our results establish a role for linker histones in maintaining centromeric chromatin integrity and proper mitotic progression in human cells. These data have implications for the regulation of transcription and higher order folding of centromeres in normal human cells; and point to potential centromere dysfunction in brain malignancies in which H1.5 is frequently highly over-expressed.

### Limitations of the study

In this study, we demonstrate that linker histone variant H1.5 plays a critical and non-redundant role at centromeric chromatin, influencing centromeric transcription and mitotic fidelity. We show that H1.5 biochemically associates with CENP-A nucleosomes, and that its depletion disrupts centromeric transcription, impairs new CENP-A loading, and leads to mitotic abnormalities.

However, certain limitations lend caution to our interpretations. First, we cannot perform genome-wide ChIP-seq profiling of all H1 variants, so the possibility of additional H1 variants contributing to centromere function remains a future question to address. Second, while our data suggest that H1.5 binding to CENP-A nucleosomes differs from canonical H1-dyad interactions, ongoing high-resolution structural studies of stable interactions between CENP-A nucleosomes and linker histones will help shed light on the structural mode of binding. We also did not directly assess RNAPII occupancy or active histone marks at the CENP-A gene locus; this remains an area for future exploration. Overall, these limitations do not detract from our main conclusion that H1.5 plays a functional role in maintenance of centromeric chromatin integrity and mitotic progression.

## DATA AVAILABILITY

The RNA-seq and ChIP-seq sequence data and additional experimental detail can be found in the Gene Expression Omnibus (GOE) under the accession number GSE242112. H1.5 ChIP-seq data used in this study was obtained from GEO with the accession number GSE166645.

## SUPPLEMENTARY DATA

Supplementary Data are available at NAR online.

## AUTHOR CONTRIBUTIONS

AS, MB, and YD designed the biological study; AS and MB performed all biochemical, cytological, and genetic experiments with assistance from RB; AS performed *in vitro* reconstitution of mononucleosomes and DM performed all AFM imaging, measurements, and analysis; AS and GA performed qRT-PCR. SB and AS performed sequencing analysis. AS, MB, and YD wrote the manuscript. All authors read and approved the final manuscript.

## ACKNOWLEDGEMENTS

The authors thank the late Dr. Craig Mizzen and the department of cell and developmental biology at University of Illinois at Urbana Champaign (UIUC), for the gift of the H1.5 antibody used in this manuscript. We thank Dr. Arthur Skoultchi for his useful suggestions and the use of the HPLC equipment in generating supporting data. We thank Dr. Stefan Dimitrov (University of Grenoble), Dr. Stephan Diekmann (retired, FLI-Jena), and Dr. Yuri Lyubchenko (University of Nebraska) for thoughtful comments on H1 and centromeres throughout the course of this work.

---

**Figure S1.**
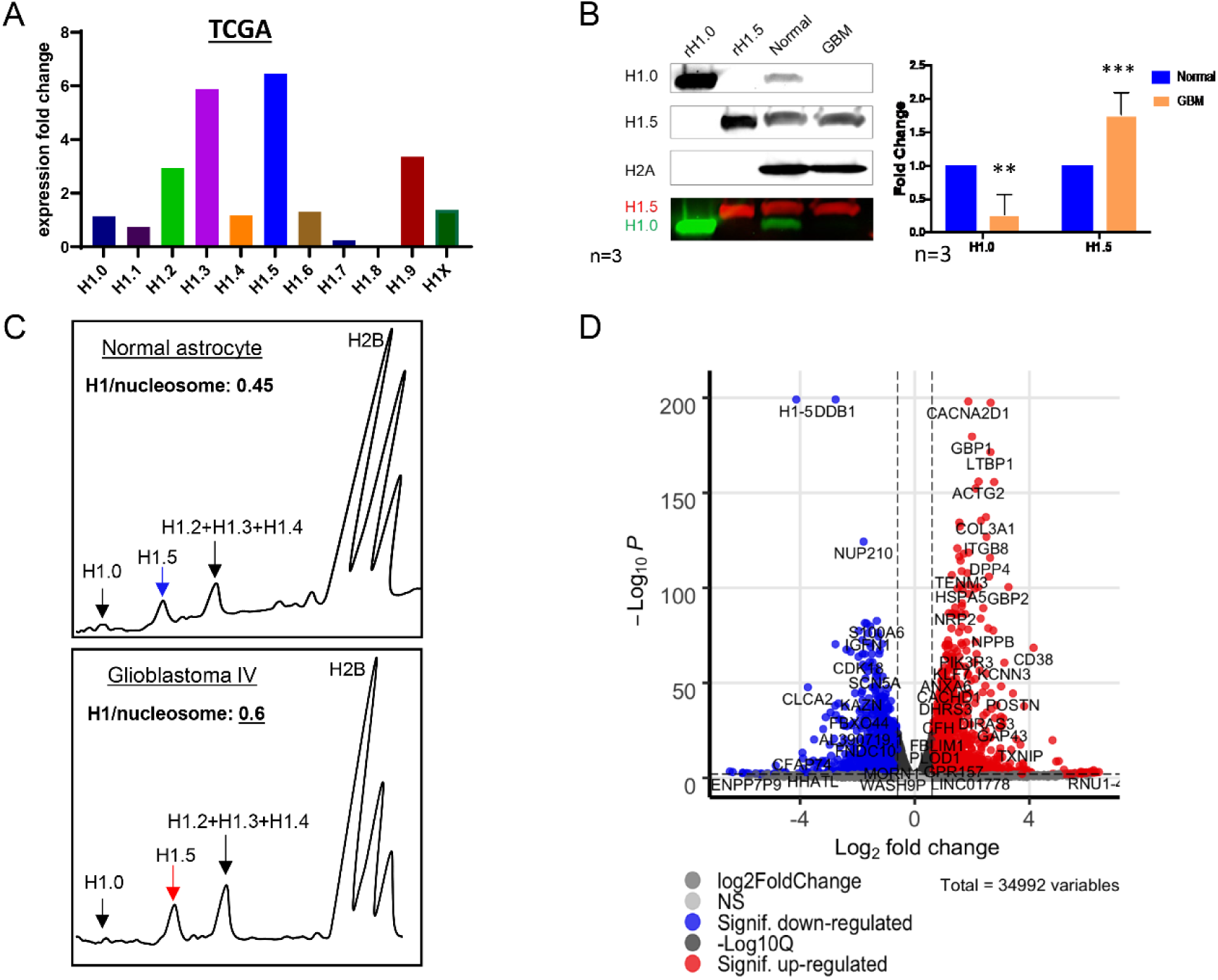
Histone H1.5 is a predominant H1 variant in astrocyte-lineage cells and regulates alpha-satellite transcription. (A) mRNA expression levels of histone H1 subtypes in Glioblastoma Multiforme (GBM) patient samples were analyzed using The Cancer Genome Atlas (TCGA) dataset. Among all variants, H1.5 shows the highest expression, suggesting a potential subtype bias in astrocytic tumors. (B) Western blot of hydroxylapatite-purified histone extracts from normal fetal astrocyte cells (SVGp12) and GBM cells (U138) confirm that H1.5 is elevated in GBM cells, accompanied by a notable reduction in H1.0. Quantification of band intensities (right) supports this subtype switch. Data are shown as mean ± standard deviation (SD) from n = 3 independent biological replicates. Statistical significance was determined using two-tailed Student’s t-test; *p < 0.05, **p < 0.01, ***p < 0.001, ****p < 0.0001. (C) High-performance liquid chromatography (HPLC) analysis of acid-extracted histones reveals that H1.5 contributes significantly to the total H1 pool in glioblastoma cells compared to normal astrocytes. Peak identities are based on known elution profiles for individual H1 subtypes. (D) Volcano plot from RNA-seq analysis of SVGp12 cells treated with siScramble versus siH1.5 identifies significantly differentially expressed genes (FDR < 0.05). Alpha-satellite transcripts and multiple mitotic regulators are among the most downregulated genes upon H1.5 depletion.

**Figure S2.**
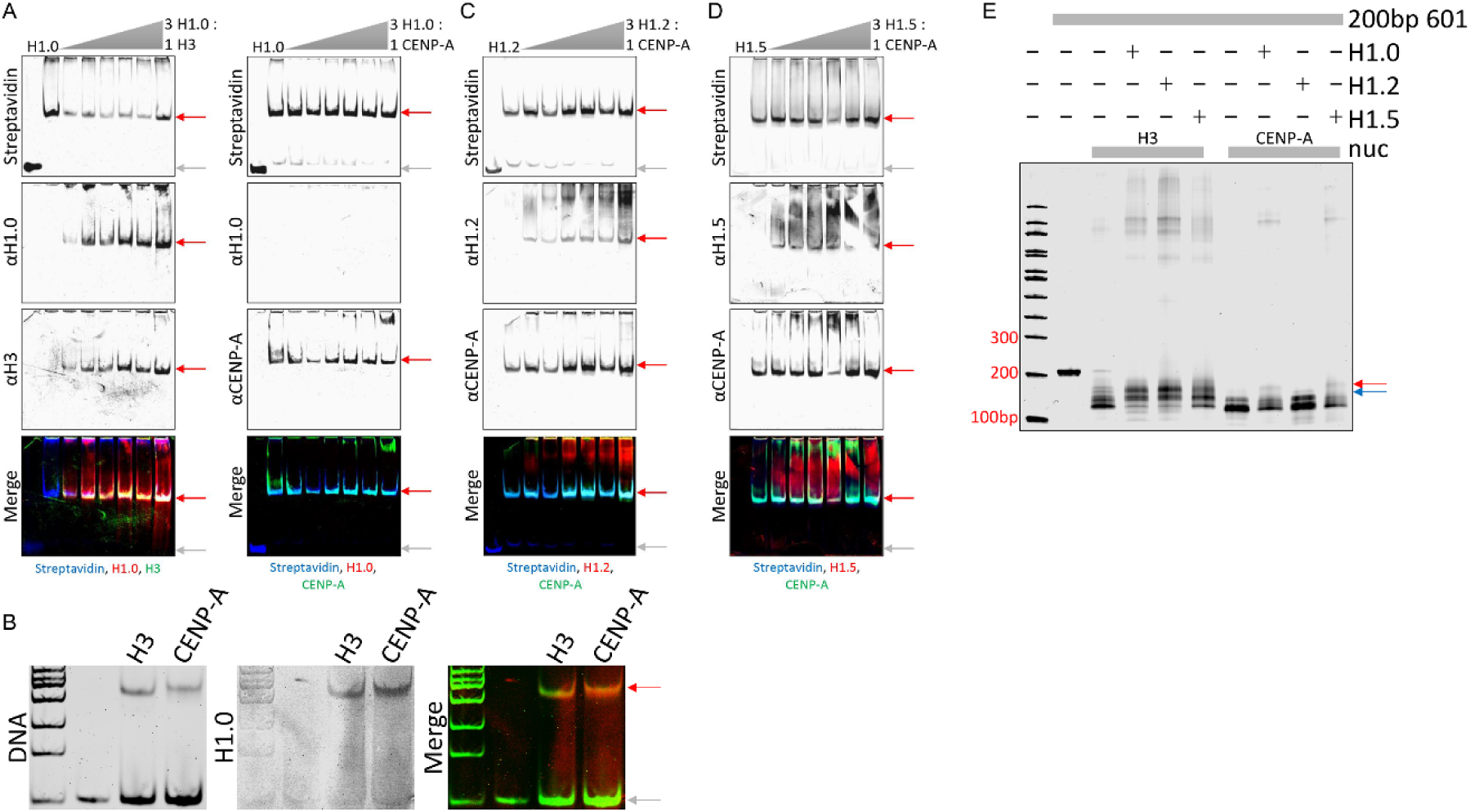
Histone H1s interact with H3 and CENP-A nucleosomes differently. (A) Histone H1.0 forms a stable nucleosome complex with H3- but not CENP-A- containing nucleosomes in the presence of 200 bp 601 positioning sequence with 5’ biotinylated ends. (B) H1.0 forms a stable nucleosome complex with both H3- and CENP-A- containing nucleosomes when using unmodified 200 bp 601 positioning sequence (no biotinylated ends). (C) H1.2 forms a stable nucleosome complex with CENP-A. (D) H1.5 forms a stable nucleosome complex with CENP-A. (E) Linker histones H1.0, H1.2, and H1.5 MNase protection signatures vary when bound to H3- or CENP-A nucleosomes.

**Figure S3.**
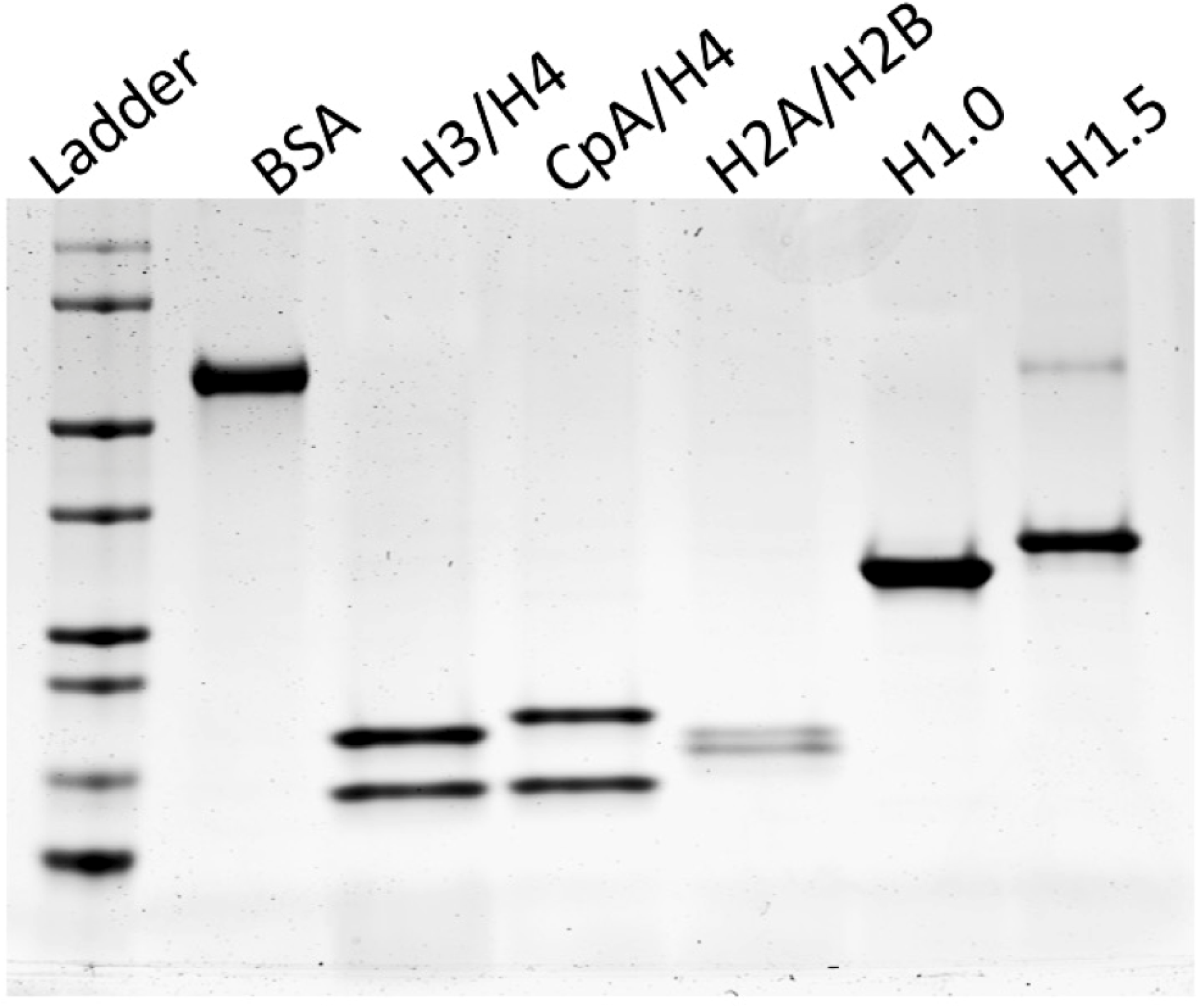
Protein validation for *in vitro* nucleosome reconstitution. Coomassie-stained SDS-PAGE gel showing purified recombinant proteins used for nucleosome reconstitution: H3/H4 and CENP-A/H4 tetramers, H2A/H2B dimers, and linker histones H1.0 and H1.5.

**Figure S4.**
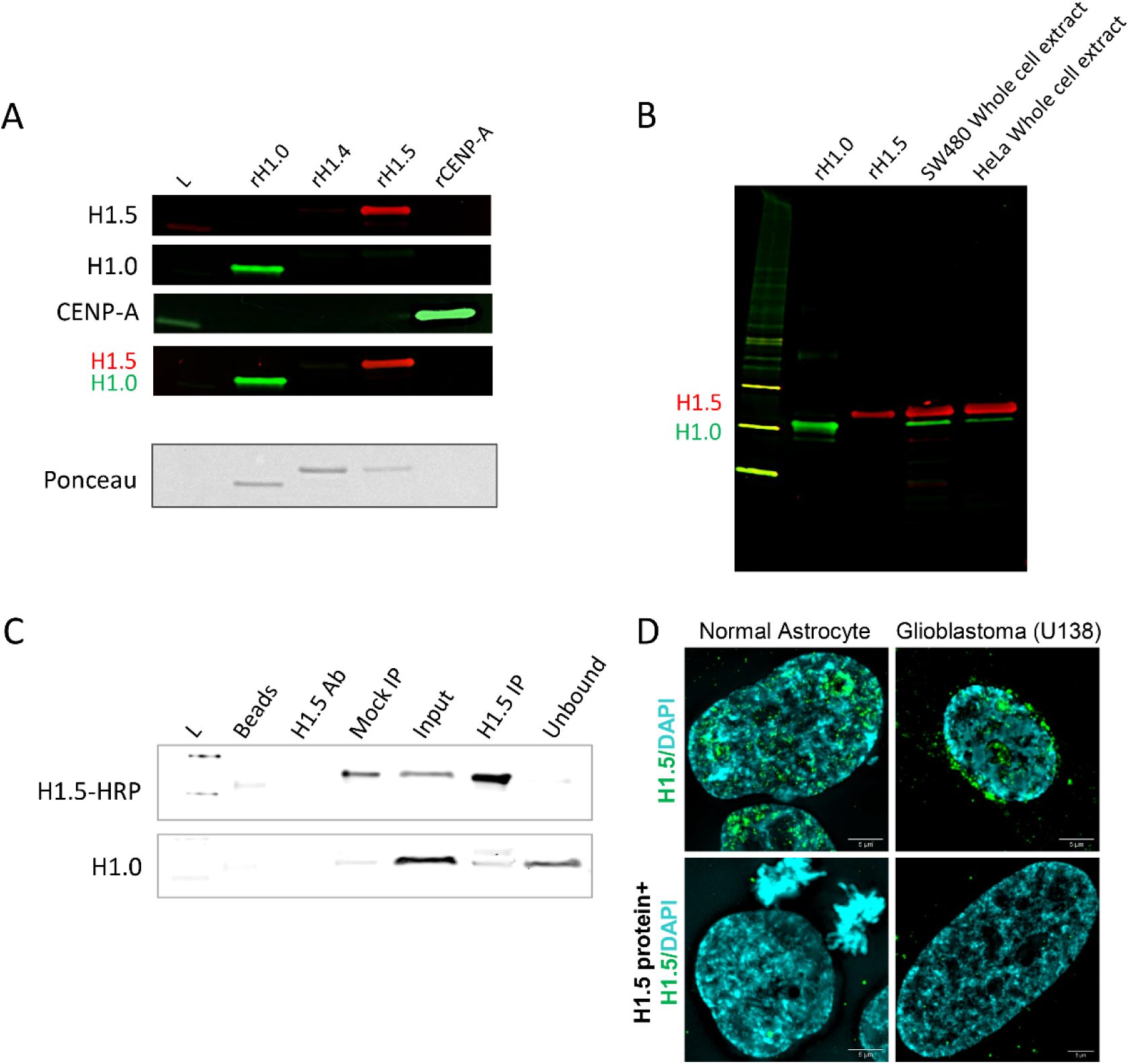
Validation of affinity-purified custom rabbit anti-H1.5 antibody for Western blotting, immunoprecipitation, and immunofluorescence. (A) Western blot showing antibody specificity against recombinant proteins H1.0, H1.4, H1.5, and CENP-A. The anti-H1.5 antibody detects only the H1.5 band (red), with no cross-reactivity to H1.0 or other tested proteins. Ponceau staining indicates equal loading. (B) Validation of H1.5 antibody specificity using histone extracts from colorectal and cervical cancer cell lines. Dual-color detection (H1.5 in red, H1.0 in green) confirms selective recognition of H1.5 across biological sources. (C) Immunoprecipitation (IP) of endogenous H1.5 from cell lysates using the custom antibody. Western blot probed with H1.5-HRP conjugate confirms efficient and specific pulldown of H1.5 compared to control IgG or H1.0 antibody IP. (D) Immunofluorescence validation of H1.5 signal in SVGp12 normal astrocytes and glioblastoma U138 cells. Pre-incubation of antibody with recombinant H1.5 protein (bottom panels) results in loss of nuclear signal, confirming specificity of detection. Nuclear morphology and H1.5 distribution differ between normal and cancer cells.

**Figure S5.**
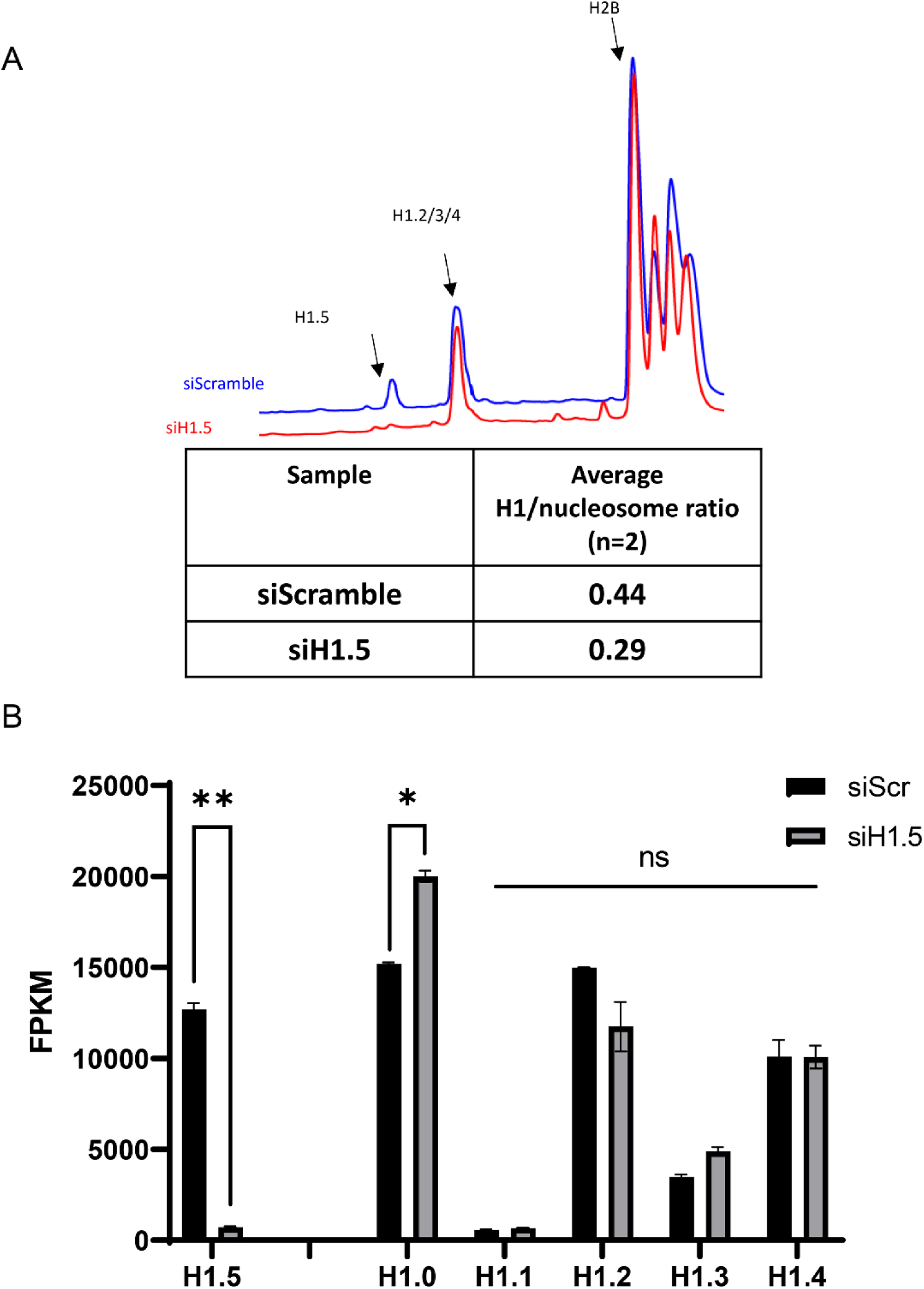
H1.5 is selectively reduced upon siRNA knockdown. (A) HPLC chromatograms of acid-extracted histones from siScramble (blue) and siH1.5-treated (red) SVGp12 cells show a selective reduction in H1.5 peak intensity following knockdown. Quantification of H1/nucleosome ratios (normalized to H2B) confirms a significant reduction of total H1 in siH1.5-treated samples (0.29) compared to siScramble control (0.44). (B) RNA-seq quantification of histone H1 mRNA subtypes reveals a significant reduction in H1.5 (p < 0.01) and a mild but significant increase in H1.0 (*p < 0.05*) upon siH1.5 knockdown. No significant change (ns) was observed in H1.1, H1.2, H1.3, or H1.4 transcripts (n = 2 biological replicates, error bars indicate SEM).

**Figure S6.**
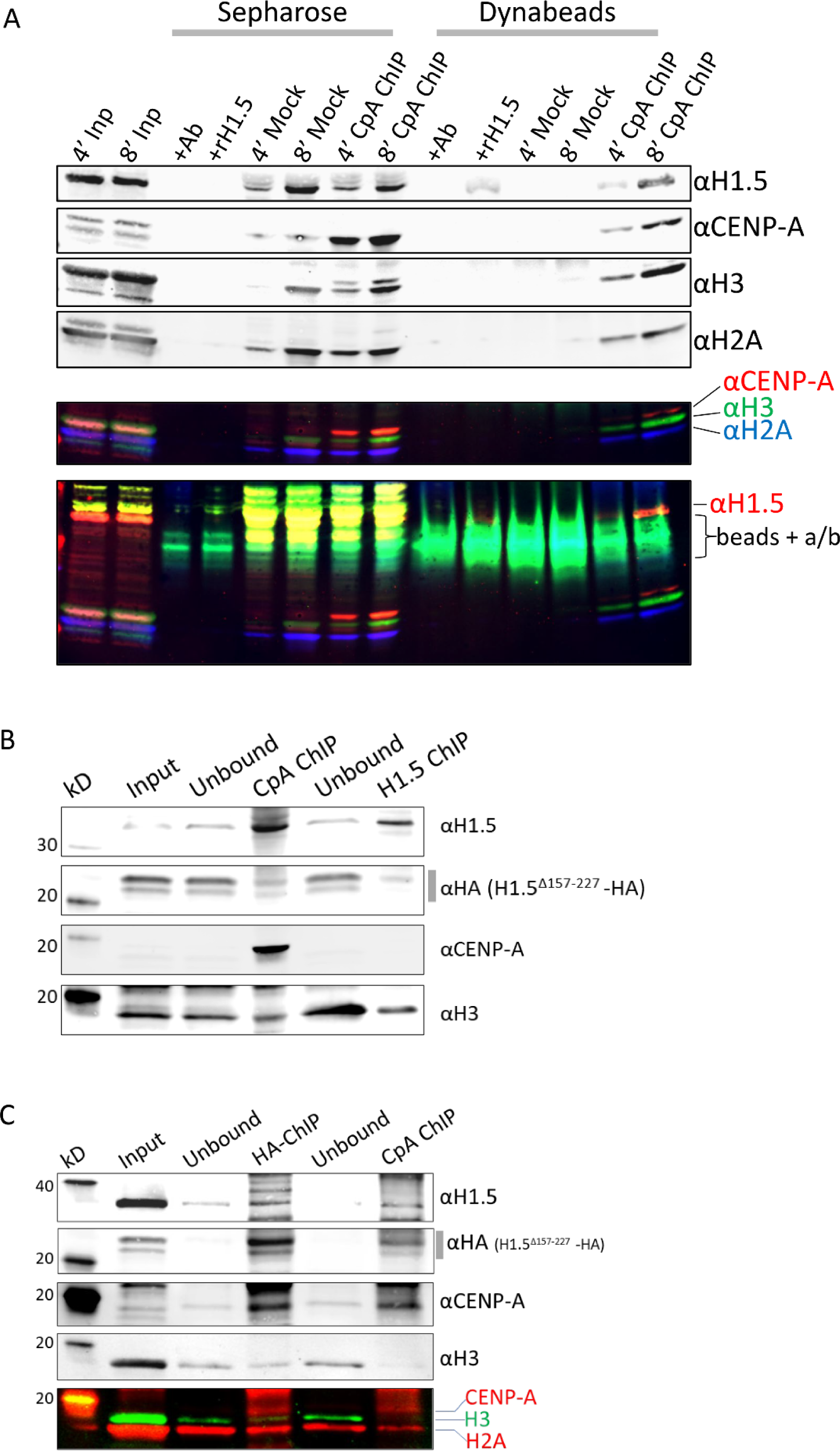
H1.5 associates with CENP-A nucleosomes in vivo. (A) In vivo ChIP verification of chromatin prepared using medium (4 min MNase digestion) or mononucleosome (8 min MNase digestion) conditions, using two bead formats: Sepharose and Dynabeads Protein A. Immunoprecipitation was performed with antibodies against H1.5, CENP-A, or control IgG. Western blotting reveals co-precipitation of H1.5 and CENP-A from chromatin, along with core histones H3 and H2A. Multiplex fluorescent detection of histone bands is shown below the Western panels. Notably, Dynabeads Protein A showed minimal non-specific binding in mock immunoprecipitated samples, and was thus used for all subsequent H1.5 ChIP experiments. (B) Co-IP using anti-CENP-A and anti-H1.5 antibodies from HeLa cells expressing wild-type or truncated H1.5 (Δ157–227-HA). CENP-A robustly co-precipitates with both endogenous and truncated H1.5, suggesting that the C-terminal tail is dispensable for the interaction. (C) Comparative Western blot analysis and fluorescent imaging of histone content in HA-ChIP and CENP-A-ChIP samples confirms the shared presence of CENP-A, H3, and H2A in nucleosome fractions pulled down with H1.5.

**Figure S7.**
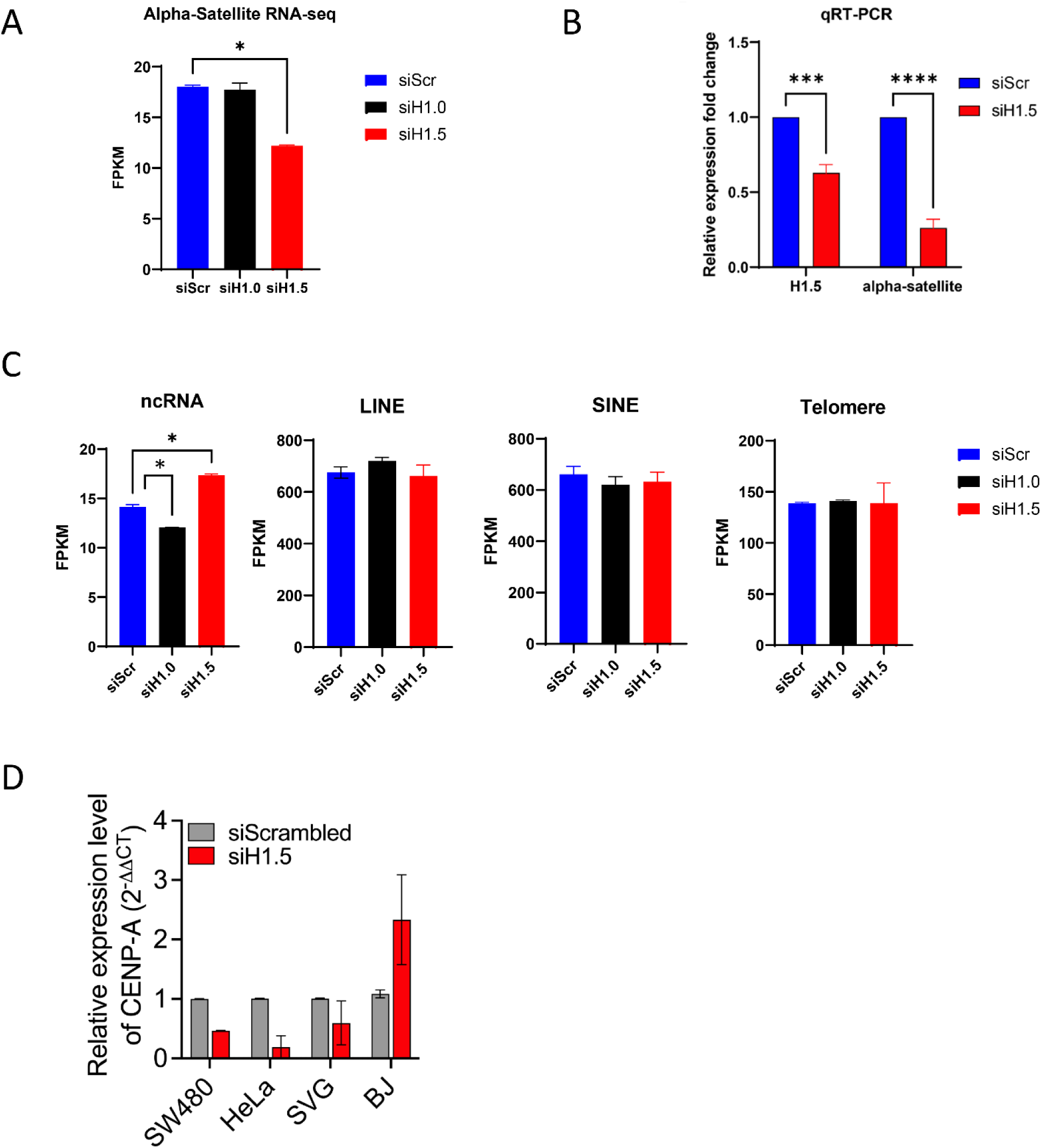
H1.5 depletion reduces α-satellite expression (A) RNA-seq quantification of α-satellite RNA in astrocytes treated with siScramble (blue), siH1.0 (black), or siH1.5 (red). H1.5 knockdown leads to a significant reduction in α-satellite expression, while H1.0 has no significant effect (unpaired two-tailed t-test, *p < 0.05) (B) qRT-PCR validation confirms effective knockdown of H1.5 and associated reduction in alpha-satellite RNA in independent biological replicates. Expression levels are normalized to control housekeeping genes and presented as mean ± SEM (p < 0.001). (C) RNA-seq analysis of additional noncoding RNA categories, including total ncRNA, LINEs, SINEs, and telomeric repeats. Only the total ncRNA pool shows a significant reduction following H1.5 depletion (*p < 0.05). (D) CENP-A mRNA levels in SW480, HeLa, SVGp12, and BJ cells after siRNA-mediated knockdown of H1.5 (siH1.5, red) and compared them to siScramble control (gray) using quantitative RT-PCR. The relative expression values are based on calculations from the 2^−ΔΔCt^ method with normalization against GAPDH. The SW480, HeLa, and SVGp12 cells show decreased CENP-A transcripts when H1.5 levels decrease because centromere transcription becomes impaired. BJ fibroblasts demonstrate elevated CENP-A expression levels which indicate potential cell-type-specific differences in H1.5 function or the existence of compensatory mechanisms. The standard deviation is shown with error bars from three biological replicate measurements.

**Figure S8.**
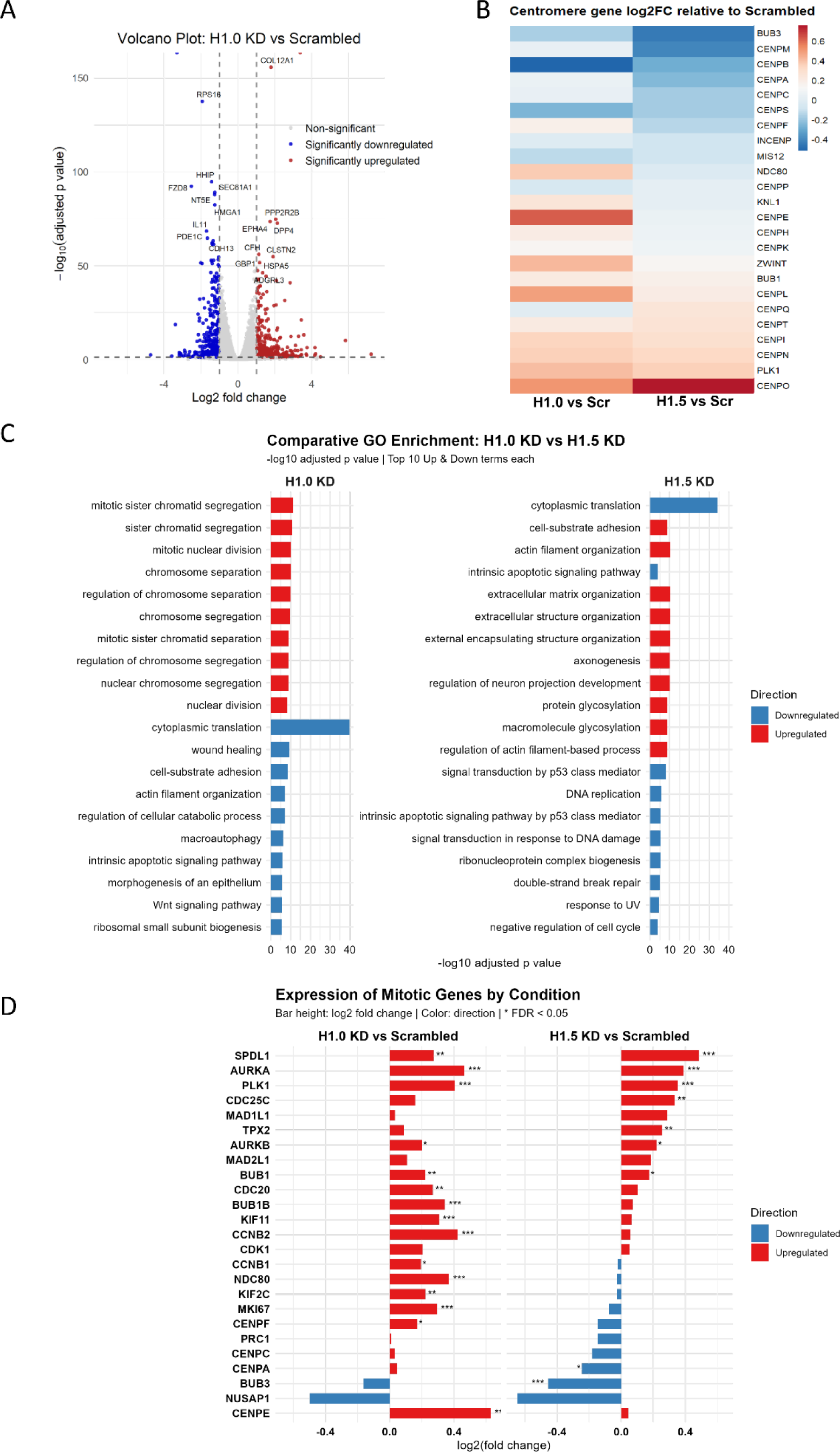
Comparative RNA-seq analysis of H1.0 and H1.5 depletion reveals distinct transcriptional effects. (A) Volcano plot showing differential gene expression upon H1.0 knockdown. Each point represents a gene, plotted by log₂ fold change (x-axis) and −log₁₀ p-value (y-axis). Genes with p < 0.05 and log₂ fold change > 1 are shown in red (significantly upregulated), and those with p < 0.05 and log₂ fold change < −1 are shown in blue, significantly downregulated). Non-significant genes are shown in light gray. Vertical dashed lines mark log₂ fold change thresholds (±1), and the horizontal dashed line indicates the p-value cutoff of 0.05. The top 10 most statistically significant up- and downregulated genes are labelled. (B) Heatmap showing log2 fold changes of curated centromere-associated genes between siH1.5 and siScramble conditions. (C) Comparative GO term enrichment analysis of the top 10 most significantly upregulated and downregulated biological processes following siH1.0 (left) or siH1.5 (right) knockdown relative to scramble control. Gene ontology enrichment was performed using clusterProfiler, with pathways ranked by adjusted p-value. (D) Targeted expression analysis of mitotic regulatory genes selected from the Reactome “M Phase” and “Mitotic Metaphase and Anaphase” gene sets (MSigDB). Bars represent log2 fold change for siH1.0 vs scramble (left) and siH1.5 vs scramble (right). Asterisks indicate statistical significance (* FDR < 0.05, ** FDR < 0.01, *** FDR < 0.001, **** FDR < 0.0001).

**Figure S9.**
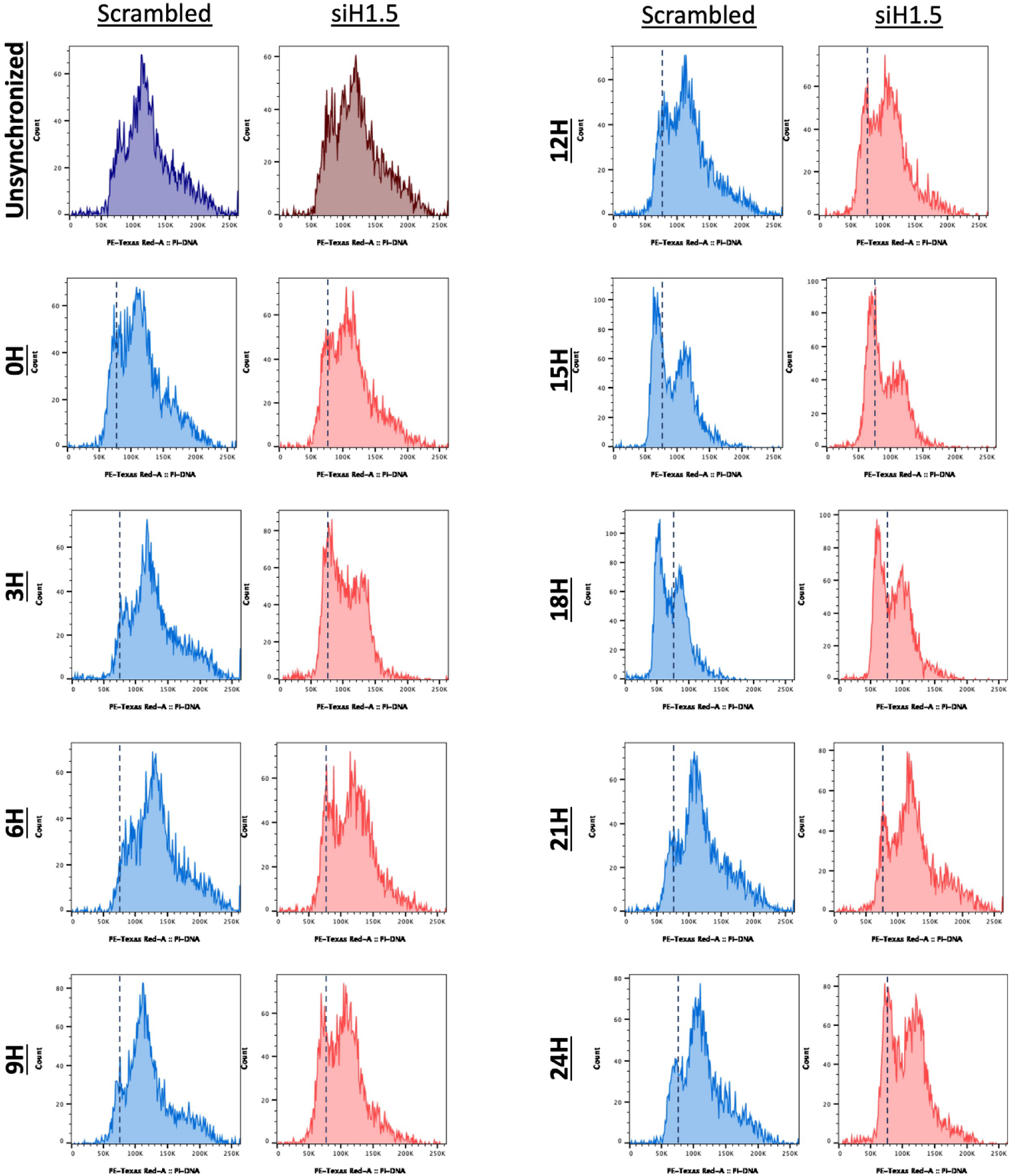
SVGp12 cells exhibit disrupted cell cycle progression after H1.5 knockdown. The representative flow cytometry (FACS) plots show propidium iodide–stained SVGp12 cells in two different conditions which include siScramble control (left panels in blue) and siH1.5-treated (right panels in red) during the cell cycle re-entry time course. A double thymidine block was used to synchronize cells before transferring them into fresh media. Cell cycle dynamics were analysed by collecting samples at specified time intervals from 0 to 24 hours post-release to measure DNA content. The DNA content analysis of siScramble-treated cells demonstrates a synchronized progression through the cell cycle from G1 to S to G2/M phases before returning to G1 within 21–24 hours. Cells treated with siH1.5 experience delayed progression through the cell cycle with significant G2/M phase accumulation and expanded peaks beginning at 9 hours after release. The observed pattern reveals a compromised mitotic exit which can be explained by the triggering of the spindle assembly checkpoint or a delayed mitosis. The top row asynchronous samples serve as reference profiles for analysis. The data demonstrates that H1.5 plays an essential role in both timely mitotic progression and successful cell division completion in human glial SVGp12 cells.

**Figure S10.**
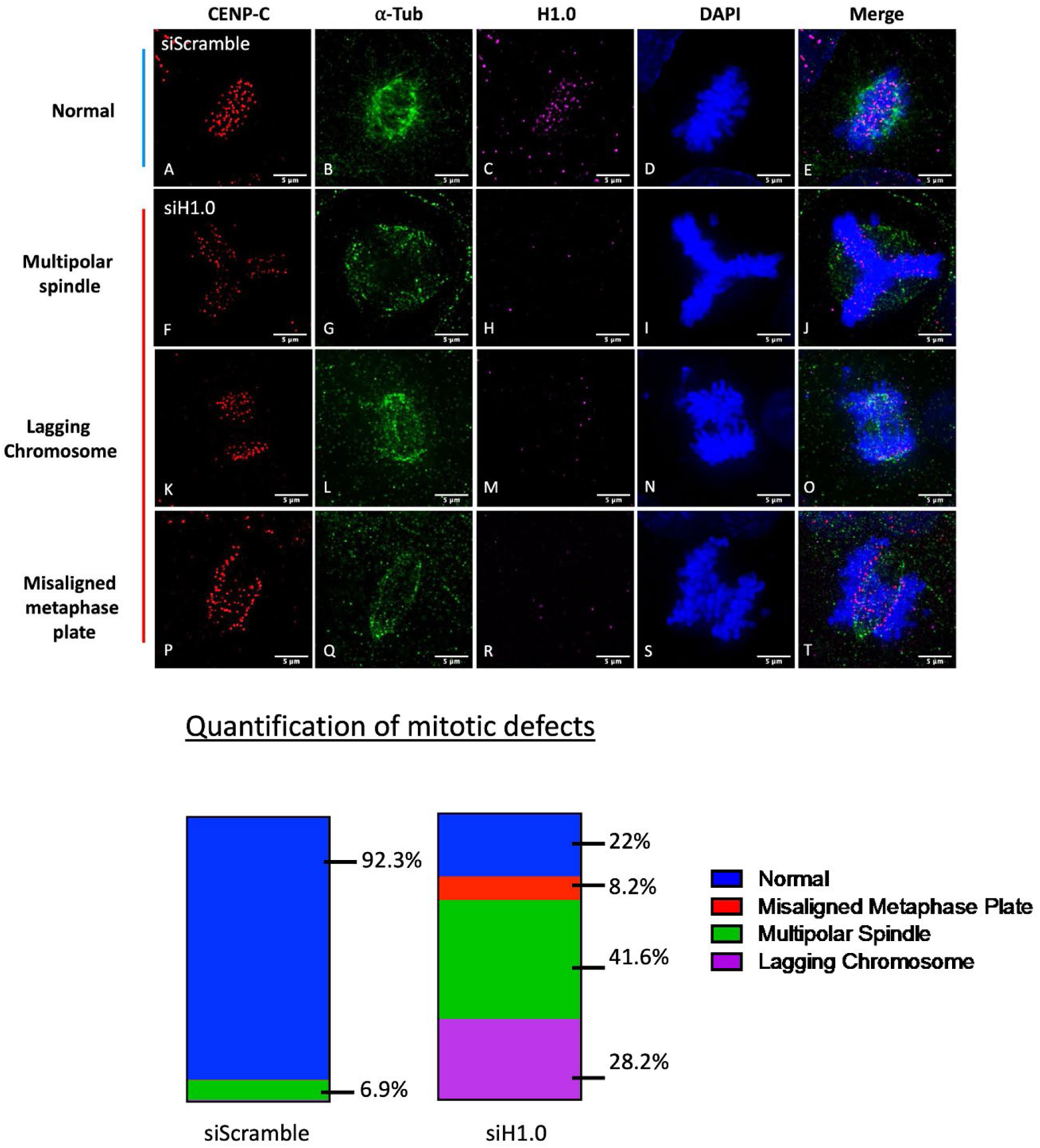
Human cells exhibit specific mitotic defects when histone H1.0 levels are reduced. SVGp12 cells transfected with either siScramble (top row) or siH1.0 (bottom three rows) show representative immunofluorescence images stained for CENP-C (red), α-tubulin (green), H1.0 (magenta), and DNA using DAPI (blue). siScramble control cells demonstrate typical mitotic progression with properly aligned metaphase plates and bipolar spindles. Cells with depleted H1.0 levels show several mitotic abnormalities which manifest as multipolar spindles (G–J), lagging chromosomes (K–O), and misaligned metaphase plates (P–T). The reduction of H1.0 signal in knockdown cells demonstrates effective siRNA treatment. The combined and separate imaging channels demonstrate the spindle organization as well as the positions of chromosomes and the placement of histones. The bottom panel contains numerical data from 200 mitotic cells analysed under each experimental condition. Analysis of siScramble-treated cells reveals normal mitotic processes in 92.3% of cases while defects occur in 6.9%. Cells with siH1.0 knockdown show significant mitotic defects where 41.6% formed multipolar spindles while lagging chromosomes appeared in 28.2% of cells and 8.2% had misaligned metaphase plates. Untreated cells with siH1.0 show that only 22% complete mitosis without any visible defects.

